# Fiberbots: Robotic fibers for high-precision minimally invasive surgery

**DOI:** 10.1101/2023.02.16.528823

**Authors:** Mohamed E. M. K. Abdelaziz, Jinshi Zhao, Bruno Gil Rosa, Hyun-Taek Lee, Daniel Simon, Khushi Vyas, Bing Li, Hanifa Koguna, Yue Li, Ali Anil Demircali, Huseyin Uvet, Gulsum Gencoglan, Arzu Akcay, Mohamed Elriedy, James Kinross, Ranan Dasgupta, Zoltan Takats, Eric Yeatman, Guang-Zhong Yang, Burak Temelkuran

## Abstract

Technologies that rely on the fundamental principle of thermal expansion have demonstrated high-precision, a growing demand in fields driven by miniaturization. However, scalable production of high aspect ratio devices that harness this capability while facilitating flexibility in design and functionality remains a challenge. We employed the high-throughput fiber thermal drawing technique to readily fabricate multimaterial fiberbots that can precisely and omnidirectionally move by asymmetric thermal expansion. These millimeter-scale fibers (< 2 mm) show excellent repeatability and linearity, negligible hysteresis, and can achieve micron-level resolution over four orders of magnitude motion range. By integrating these robotic fibers with medical devices that can perform cellular-level tissue imaging, diagnosis, and manipulation, we showcase their versatility through benchtop and preclinical animal studies and their overall potential impact on medicine, biomedical engineering, robotics, and beyond.

**One Sentence Summary:** Scalable manufacturing and integration of robotic fibers that deliver high-precision motion when heated.

## INTRODUCTION

Materials tend to expand as temperatures increase sometimes leading to undesired consequences such as bent rail tracks on hot days and malfunctioning electronics. However, positive and negative thermal expansion can also be usefully harnessed in different ways and scales, such as in mercury thermometers for health status assessment, miniature circuit breakers for electrical protection, optical fiber cantilevers, and microelectromechanical systems (MEMS) for micro- and nano-manipulation *(1 - 3)*, and more recently, soft actuation devices for untethered robots *(4, 5)* and artificial muscle control *(6 - 9)*. Although actuators relying on this intrinsic material property have shown great potential in achieving precise displacements and large forces, flexibility in material and design choices coupled with the ability to integrate other functional platforms that target emerging trends such as precision surgery (e.g., microrobotic laser steering *(10)*) remains a challenge *(11)*. Moreover, in this burgeoning field of minimally invasive and robotic surgery, there is a growing interest in instruments that can be: (i) delivered to challenging surgical sites (endoluminal or endovascular) in tandem with traditional imaging modalities, (ii) utilized to maneuver other diagnostic and therapeutic instruments with high-precision and spatial resolution within the safe limits of the human body, and (iii) for the targeted delivery of pharmacological or physical therapies (e.g. ablation) by reducing the size of injury and potential infections, which could potentially lead to faster recovery times.

We attempt to address the challenging task of bringing high-precision robotic control to minimally invasive surgery by fabricating long, thin, and flexible electrothermal actuators that deliver “*in vivo*” precision using the thermal fiber drawing process. Initially developed for silica glass optical fibers, the thermal fiber drawing process has become a significant tool in the scalable fabrication of structurally flexible and intrinsically integrable devices. The first multimaterial fibers were realized by Hart et al. and Temelkuran *et al*. in 2002, which comprised 21 alternative bi-layer structures of polymer and chalcogenide glass to create a Bragg mirror inside the core of the fiber *(12, 13)*. This led to a novel way of guiding light that facilitated the transmission of wavelengths not feasible with single material (silica) fibers. In 2004, Bayindir *et al*. demonstrated for the first time a metal-insulator-semiconductor structure in a fiber that was used to fabricate a flexible photo-detection fiber that was sensitive to visible and infrared light across its length *(14)*. Since then, thermal drawing has been used to fabricate fibers for neural interfacing *(15 - 17)*, deformation sensing *(18, 19)*, electrochemical sensing *(20, 21)*, electromechanical actuation *(22)*, artificial muscles *(8)*, additive manufacturing *(23)*, soft robots *(24, 25)*, and smart textiles *(26, 27)*, among others *(28, 29)*.

The process starts from a macroscopic, single, or multimaterial preform, usually centimeters in diameter and tens of centimeters in length. Preforms can be made by various combinations of techniques, including but not limited to the consolidation of rolled thin-films *(13)*, vacuum compression molding of pellets *(30)*, machining of bars and rods, additive manufacturing *(31 - 33)*, and stack-and-draw approach (Holey fibers) *(34 - 36)*. Examples of preform materials include polymers, glass, metal alloys (amorphous, liquid, low melting point), semiconductors, and dielectrics. Once fabricated, the preform is thermally softened above its glass transition temperature (*T*_*g*_) at a draw tower and pulled in the viscous fluid state to the desired diameter (tens to thousands of micrometers) and length (tens to hundreds of meters); while maintaining the transverse geometry across the fiber length *(37)*.

Here, we describe an electrothermally actuated fiber device (referred to as fiberbot in the remainder of the text) resulting from the cross-cutting themes of fundamental physics, manufacturing technologies, medical device design, and robotic systems. This work initially lays out the actuator’s design, which encompasses a thin-walled polymeric fiber embedded with thin metallic resistive wires. The localized Joule heating of the wires generates a thermal gradient across the fiber’s cross-section, which induces an asymmetric thermal expansion and the desired high-precision motion of the fiberbots. Second, we present a rapid prototyping approach that simplifies the production of hundreds of meters of these inexpensive, bespoke, and integrable actuators that can readily function. Next, to control these actuators, we implement open-loop and closed-loop (i.e., optical and temperature-based feedback) controllers to trace pre-defined paths accurately. We further demonstrate the versatility of the proposed actuators by exploiting their tubular configuration to teleoperate miniaturized electromagnets and maneuver minimally invasive diagnostic (e.g., fiber bundles for optical biopsy) and therapeutic (e.g., surgical laser fibers) tools. In addition, we showcase the possibility of using these fiberbots to create standalone surgical tools, which we employ in standard laparoscopic interventions and integrate with commercially available robotic surgical systems. Last, in this work, we validate the efficacy and safety of the fiberbots as interventional medical devices by using them to move surgical fiber lasers with high precision in realistic *in vivo* conditions.

## RESULTS

### Scalable fabrication and design of the high-precision fiberbots

The scalable fiber drawing process was employed to draw polymeric fibers from 3D printed preforms (Fig. 1, A and B). This process facilitates the rapid and low-cost production of these high aspect ratio materials with arbitrary cross-sectional designs. Comparing different widely 3D-printable and thermally drawable thermoplastics, we identified polycarbonate (PC) as a suitable candidate for the polymeric constituent of the fibers (*T*_*g*_ = 112-113°C, coefficient of linear thermal expansion α = 2.1 × 10^−5^ °C^-1^, isotropic thermal conductivity = 0.22 W/(m.K)). In addition to its relatively high thermal expansion coefficient and low thermal conductivity, PC has biocompatible grades, high mechanical strength, and dimensional stability, making it an ideal candidate for producing medical devices. Forty-four gauge (44 AWG) stainless-steel wires (resistivity ρ = 7.4 × 10^−7^ Ω.m) were co-fed and embedded longitudinally within the fiber during fabrication (Fig. 1, C and E). The flow of electric current through these wires generates heat, leading to a localized increase in the temperature of the enveloping polymer and their combined thermal expansion (Fig. 1F). Finite element analysis was applied to verify the feasibility of the proposed actuation mechanism and optimize the fiber’s cross-sectional structure (fig. S1, fig. S2, fig. S3, and movie S1). The final design of the fiberbots comprised a hollow PC cylinder and four pairs of equidistantly spaced stainless-steel wires around the cylinder’s circumference. The four pairs of wires facilitate the thermal expansion triggered cantilever bending of the fiber in two planes. On the other hand, the hollow-core configuration provides a means to introduce other diagnostic and therapeutic tools, reduces the fiber’s overall material to be heated, and introduces a thermal barrier, thus enhancing the device’s dynamic range and response (fig. S3).

**Fig. 1.**
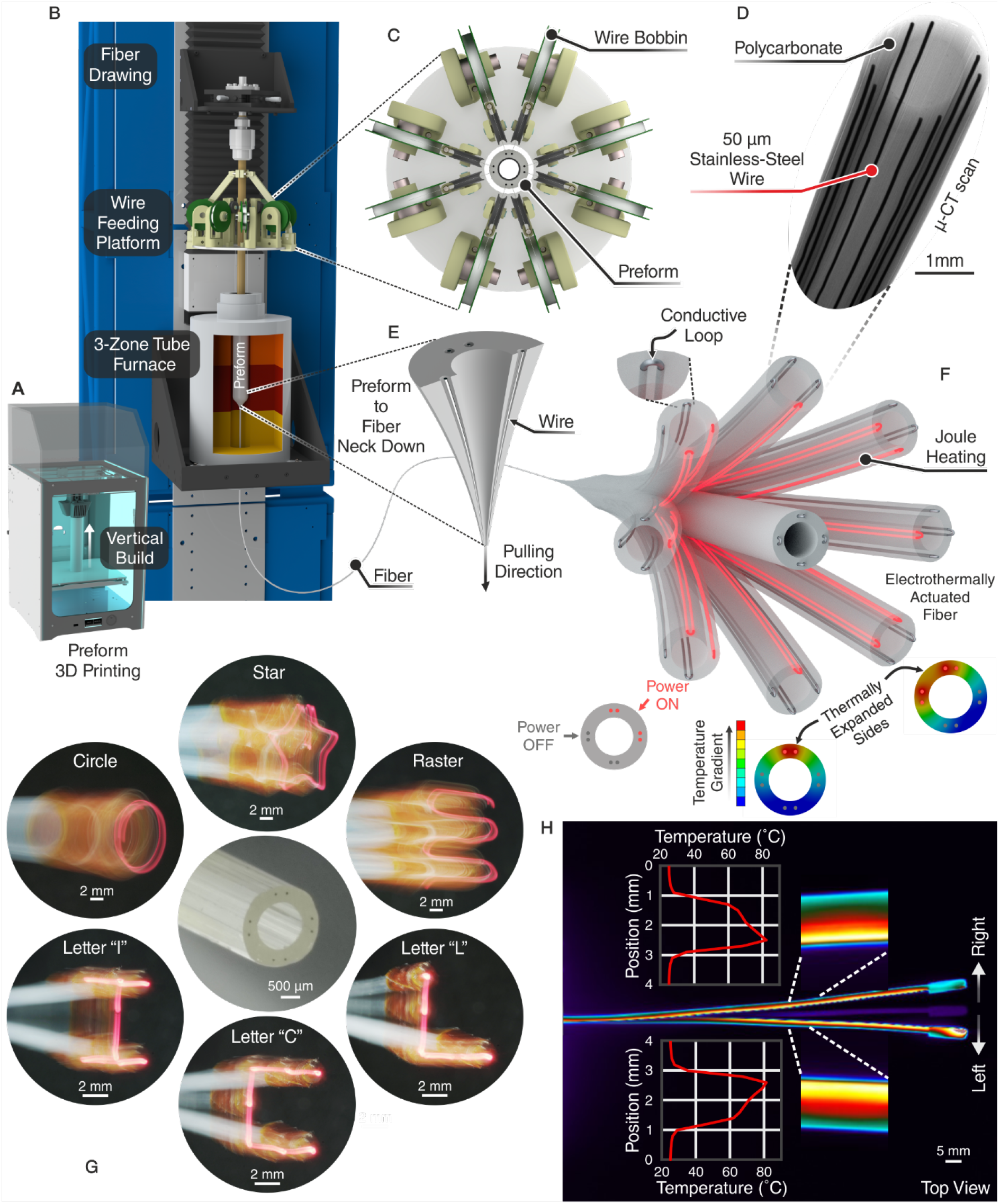
Fabrication and working principle. (**A**) Preform fabrication process via 3D printing. (**B**) Electrothermally actuated fibers produced by thermal fiber drawing. (**C**) Top view of the wire feeding platform used to guide eight stainless-steel wires into the preform’s side channels. (**D**) Micro-computed tomography (μ-CT) scan of the thermally drawn fiber. (**E**) Preform- to-fiber neck-down and the subsequent passive pulling of the wires by the encircling polymer. (**F**) Illustration of the fiber’s cantilever bending, which is powered by thermal expansion. Inset: Exposed wires at the fiber’s distal end intertwined in pairs and coated with silver paint to form a conductive loop. (**G**) Cross-sectional image of the fiber (center) and six tip patterns captured using a mirrorless camera’s slow shutter speed function while moving a 500 μm optical fiber connected to a 650 nm LED. (**H**) Temperature distribution profile along the fiber’s length obtained by a thermal camera during left and right bending movements.

The fibers were drawn from 3D printed (vertically built) preforms composed of PC (40 mm outer diameter, 24 mm inner diameter, 2 mm lateral through holes, 200 mm long) (Fig. 1, A to C). The outer diameter of the fiber was tuned between 1.5 mm and 2 mm over a 20 m length by specifying the down-feed speed (i.e., speed with which the preform is fed into the furnace, *v*_*f*_ = 1 mm/min) and draw speed (speed with which the fiber is pulled, *v*_*d*_ = 0.4 m/min to 0.7 m/min). Simultaneously, the eight resistive wires were fed through the 2 mm side channels and then passively pulled by the encircling polymer as the channels within the preform reduce in size (Fig. 1E). Post-drawing, the fibers were cut to desired lengths (10-15 cm), and the wires were exposed at both ends by incising the fiber’s outer surface and pulling out the polymeric segment. The exposed wires at the fiber’s distal end were intertwined in pairs, and conductive silver paint was applied to bond each pair and ensure reliable electrical contact. Kapton tape was then used to isolate and insulate the four pairs. The wires at the proximal end were connected to a computer-controlled electronic circuit to control the potential difference across the four independent pairs. Figure 1H demonstrates an overlay of the thermal images of the PC fiber’s (outer diameter 1.65 mm and length 12 cm) planar bending upon the application of 12 V direct current (DC) (power 1.8 W) across a single pair of wires. In response to a temperature gradient of ΔT ≈ 20°C, developed along the fiber’s cross-section within three seconds, the tip of the fiber translated approximately 7 mm (left and right) from the equilibrium position.

### Fiberbots characterization

The fabricated fibers were further characterized in a well-controlled environment by investigating the relationships between power, displacement, fiber length, temperature difference, force, and velocity (Fig. 2). The bespoke setup included manual positioning stages for geometry alignment, a laser displacement sensor, and an optical microscope to focus on the fiber’s distal tip to acquire accurate two-dimensional deflection (position) information (Fig. 2A). From the same draw, using different draw speeds, two fibers with different outer diameters (1.65 and 2.00 mm) and similar lengths (12 and 10 cm) exhibited a linear relationship between input electrical power and fiber tip displacement (Fig. 2C). This was further supported by the linear correspondence between power and temperature gradient (Fig. 2I). As expected, the thinner 1.65 mm fiber demonstrated a larger displacement (by a factor of 2.1) for the same input power. Moreover, the behavior of the 1.21 times thicker fiber was repeatable across five runs with a pooled standard deviation of 27 μm. Using the same characterization technique of powering only a single pair of wires at a time, we obtained the corresponding power versus displacement curves for the remaining three directions of motion. As shown in Fig. 2D, we can control the steps of motion of the 2.00 mm fiber down to 30 μm over a dynamic range of 1 mm, which is still well above the noise level of the measurement setup. Additionally, our preliminary demonstration of the fiber moving along a sub-micron circular path outlines the capability of achieving higher resolution motion (movie S2). The force generated at zero displacements by the tip of the same fiber was repeatedly linear with respect to the input power and in the range of tens of millinewtons (20.7 mN at ≈ 1.153 W), i.e., up to seven times the fiber’s weight (3.1 mN) (Fig. 2E). Since the device’s length affects its response to thermal expansion, we examined this relation by mechanically clamping the fiber at different points (Fig. 2, B, G and H). The shorter (3.7 cm) free end of the 2.00 mm fiber produced a smaller displacement (by a factor of 1/6) for the same input power of ≈ 1.446 W. Thermal images were also captured to visualize better this effect on the 3, 6, and 9 cm free ends of the 1.65 mm fiber (Fig. 2H). Given that the resistance of conductive materials (stainless-steel) is a function of temperature, it was important to characterize this effect during fiber actuation. We measured both the voltage and current signals on the actuated stainless-steel wires, thereby calculating the resistance of the wires in real-time. A small resistance change of 2.8 Ω was recorded using differential amplifier circuitry (fig. S4B) and repeatable across five cycles of 10 V step inputs with an average tip displacement of 2.1 mm (Fig. 2J). To conclude the characterization of the fiberbots, we assessed the thicker fiber’s velocity profiles (starting from rest) for different input voltages steps (Fig. 2K). The velocities exponentially decreased as the tip approached the steady-state position with an average time constant of τ_avg_ = 1.4 s, and maximum achievable velocities ranging from 0.008 to 0.8 mm/s for step input voltages from 1.118 to 10 V (fig. S5A). Furthermore, the average time required by the fiber to move from 10% to 90% of its fully developed steady-state positions (i.e., rise time) was 8.6 s (fig. S5B). The fiberbots also exhibit a small hysteresis of 87 μm within the actuation range of 3.21 mm, which makes them well suited for applications (e.g., robotics) that require precision over a continuous range (Fig. 2L).

**Fig. 2.**
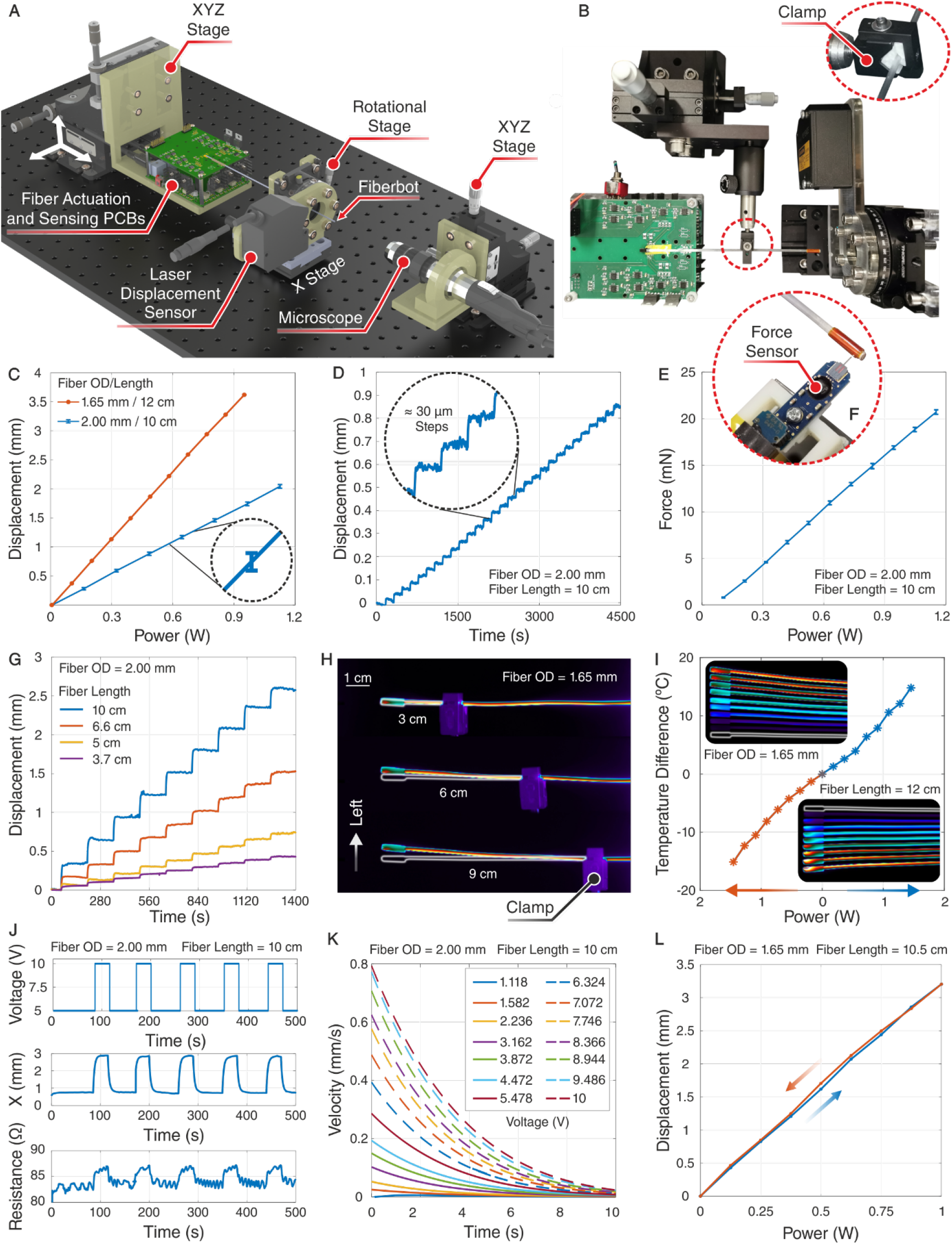
Mechanical characterization of the fiberbots. (**A**) Setup developed for characterizing the fiber within a well-controlled environment. (**B**) Extension of the setup to characterize the effect of length on the fiber’s thermal expansion and its subsequent tip displacement. (**C**) Fiber tip displacement as a function of the applied power for two fibers with different outer diameters (OD) and lengths. Inset: Error bar indicating a standard deviation (0.028 mm) obtained from five measurements using the laser displacement sensor. (**D**) Achieved positional resolution for incremental input power steps of approximately 13 mW, intermitted by 2.5 minutes. Inset: zoomed-in view of the 30 μm tip displacements. (**E**) Contact force generated laterally by the fiber tip as a function of the applied power (average of five runs). (**F**) Commercial force sensor used in the experiment. **(G)** Tip displacements for different unconstrained fiber lengths. (**H**) Top view of the thermal images of the fiber movement in the left direction during the clamp testing. (**I**) Temperature gradient measured as a function of the applied power for horizontal displacements. (**J**) Waveforms for the input voltage (top), the corresponding tip displacement (middle), and the change in resistance of the embedded stainless-steel wires (bottom). (**K**) Change in velocity of the fiber tip for different step input voltage steps (starting from rest). (**L**) Hysteresis for a fiber with an outer diameter of 1.65 mm and 10.5 cm in length, measured by performing an input power sweep from 0 W to 1 W in 125 mW incremental steps.

### Outer surface characterization

As the surface temperatures of the actuated fiberbots are above the biological safe limit (Fig. 1H), we placed the fiber alongside the laser displacement sensor inside an incubator at 38°C ± 1°C to simulate the human body and examine active cooling by injecting compressed air through the fiberbot’s central channel (Fig. 3A). Our experiments show that using air at room temperature; we can decrease the surface temperature of the actuated side by approximately 45.5°C, down to 43°C. However, the temperature gradient between the opposing ends dropped from 24.2°C to approximately 9.2°C at 5 bar, and fiber tip displacement decreased by 1.94 mm. Using a fiber that is 1.65 mm in outer diameter and 11 cm in length, the maximum fiber tip displacement to avoid thermal tissue damage is 1.68 mm for an input power of 1 W (Fig. 3B).

**Fig. 3.**
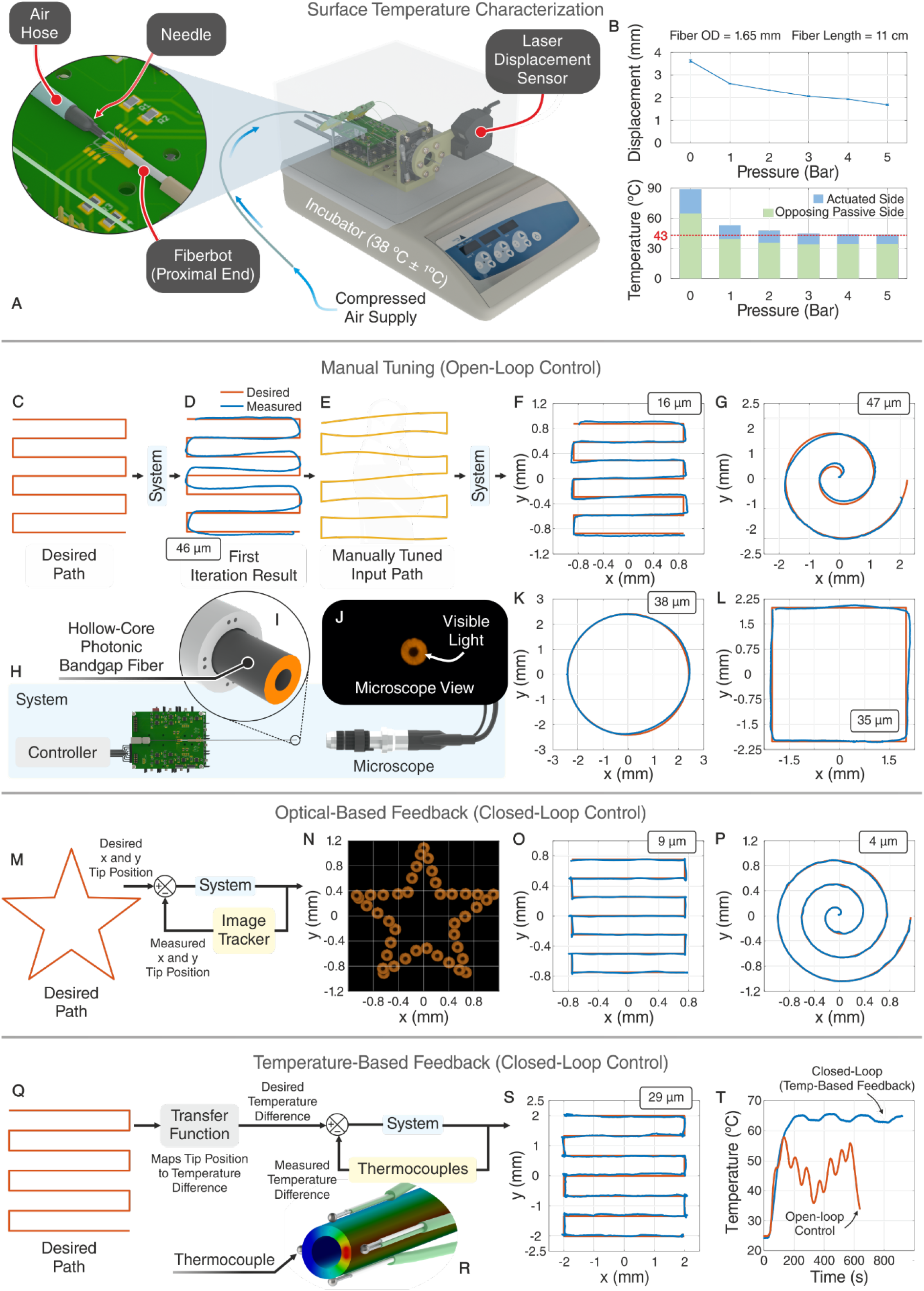
Surface temperature characterization and optimization of the fiberbot motion using different compensation mechanisms. (**A**) Setup developed for demonstrating the ability to reduce the surface temperature of the fiber upon the application of compressed air through its central channel (experiments conducted at 38°C ± 1°C environment). Inset: 21-gauge needle connected to an air hose and inserted into the central channel of the fiber. (**B**) Surface temperature of the fiber’s actuated side versus the opposing passive side at different pressures. The red dashed line indicates the biological safe limit of 43°C (top), the change in fiber tip displacement due to the cooling effect, and the reduction in the temperature gradient (bottom). (**C**) Desired raster path. (**D**) First iteration result (average path error indicated in the box) [path time: 100 s]. (**E**) Manually tuned input path based on several iterations. (**F, G, K, L**) Measured (blue) versus desired (orange) paths based on several manually tuned inputs (**F**) raster [184 s], (**G**) spiral [42 s], (**K**) circle [32 s], (**L**) square [97 s]. (**H**) System used to track the fiber tip displacements (a blue shaded box is used to represent the system in the Figure). The controller maps the desired fiber tip displacement mapped into four independently controlled and timely tuned voltage inputs across the pairs of wires. (**I**) Embedded 900 μm hollow-core photonic bandgap fiber connected to a 650 nm LED for tracking purposes. (**J**) Microscope view of the embedded fiber. (**M**) Block diagram of the closed-loop controller based on vision-based feedback. (**N**) Overlay of the acquired microscope images for a star path. (**O, P**) Measured (blue) versus desired (orange) paths based on the vision-based feedback approach [raster: 244 s, spiral: 641 s]. (**Q**) Block diagram of the closed-loop controller based on temperature-based feedback. (**R**) Illustration of four miniature thermocouples attached to the outer surface of the fiber next to the four pairs of wires to provide real-time temperature measurements. (**S**) Measured (blue) versus desired (orange) paths based on the temperature-based feedback approach [737 s]. **(T)** Average temperature detected during fiber actuation using open and closed-loop controllers (with temperature feedback) for an equivalent path.

### Open-loop and closed-loop position control

Post-characterization, the fiberbots were tested by commanding them to follow several pre-defined desired paths (Fig. 3 (D to F)). The paths comprised a set of points on the projected orthogonal plane, which are mapped into four independently controlled and timely-tuned voltage inputs for the pairs of wires (utilizing the measured relation between power and displacement). The mapped voltages were then sequentially sent from the computer to the fiberbot’s electronic interface. Next, we embedded a hollow-core photonic bandgap (PBG) fiber (0.9 mm outer diameter) coupled to a 650 nm light-emitting diode and used the optical microscope alongside a pattern matching algorithm to track the PBG fiber tip (Fig. 3, H, I and J). As shown in Figure 3D, the measured path deviates from the desired path, with an average path error (i.e., the time-independent difference between the desired and measured paths) of 45.5 μm. The inaccuracies were primarily due to the actuator’s large time constant, caused by the polymer’s intrinsically low thermal conductivity. This delay causes the fiberbots to move toward the current command’s desired position without reaching the previous command’s desired target. Another contributing factor is gravity, which becomes more significant due to the decrease in the polymer’s Young’s modulus as its temperature rises and the alignment of its molecules relaxes. These effects were compensated for by: (a) slowing down the speed with which new desired positions are reached in areas with sudden changes in direction, (b) manually tuning the magnitude of the input voltages based on previous trials (fig. S6 (D to K), Fig. 3E). Using this iterative compensation technique, we decreased the average path error of a raster by 65.9% (Fig. 3F) and similarly optimized the fiberbot’s performance for spiral (Fig. 3G), circle (Fig. 3K), and square (Fig. 3L) paths. Although encouraging, manual tuning is a time-consuming and path-dependent process, which prompted us to explore controlling the fiberbot by automatically regulating its tip position using feedback-based control. This was initially achieved by employing an optical-based approach, where we utilized the real-time tracked tip position (system’s output) to form a closed-loop system, alongside a proportional–integral–derivative (PID) controller (Fig. 3O, Fig. 3P, fig. S7A). Figures 3O and 3P demonstrate the raster and spiral paths followed by the fiberbot with average path errors less than 10 μm. Next, we investigated an alternative practical feedback approach, where we attached four miniature thermocouples to the outer surface of the fiber next to the four pairs of wires (Fig. 3R). We mapped the desired fiber tip displacement to the corresponding temperature difference between the opposing sides of the fiber using the previously characterized linear relationships between power, displacement, and temperature difference (Fig. 2, C and I). The desired temperature differences were subsequently used as setpoints for the temperature-based feedback PID controller (Fig. 3Q, fig. S7B). For a 4 by 4 mm raster path, the fiber could follow the desired path with an average path error of 29 μm (Fig. 3S). The average temperature measured along an equivalent raster path for the open and closed-loop controllers reveals a smoother evolution of the temperature profile over time for this latter mechanism, though requiring more time to complete the path due to the slow dynamics of temperature measurement (Fig. 3T).

### Magnetic tele-micromanipulation

To illustrate the potential applications of the fiberbots, we designed a vertically oriented micro-manipulation system, where we attached an electromagnetic coil (2.5 mm outer diameter) to the tip of the fiberbot. The fiberbot-coil assembly was employed to telemanipulate a small magnetic object (immersed in a water-filled Petri dish) through different paths and tasks (e.g., maze, Fig. 4 (A to E), and movie S4). Unlike the previous characterizations and position control experiments, where the fiberbot followed a pre-defined path, herein, the fiberbot was teleoperated by a user using a joystick controller (Fig. 4D, fig. S8). A microscope was placed at the bottom of the vertical setup (below the Petri dish) for visual feedback, whereas an auxiliary switch was used to pick (coil on) and drop (coil off) the particle, respectively. Figures 4C and 4D show a maze printed on transparent paper attached to the Petri dish and the microscope view of the maze pattern with a total area of 5.2 × 7.6 mm^2^ and a narrowest wall gap of 600 μm. Using the same simulated environment, the fiberbot was commanded to pick and place the object in different targets (movie S4), weave through football (soccer) cones (movie S5), and pass-through defenders to score a goal (movie S5). These demonstrations could potentially inspire applications which demand interaction with small particles in confined spaces.

**Fig. 4.**
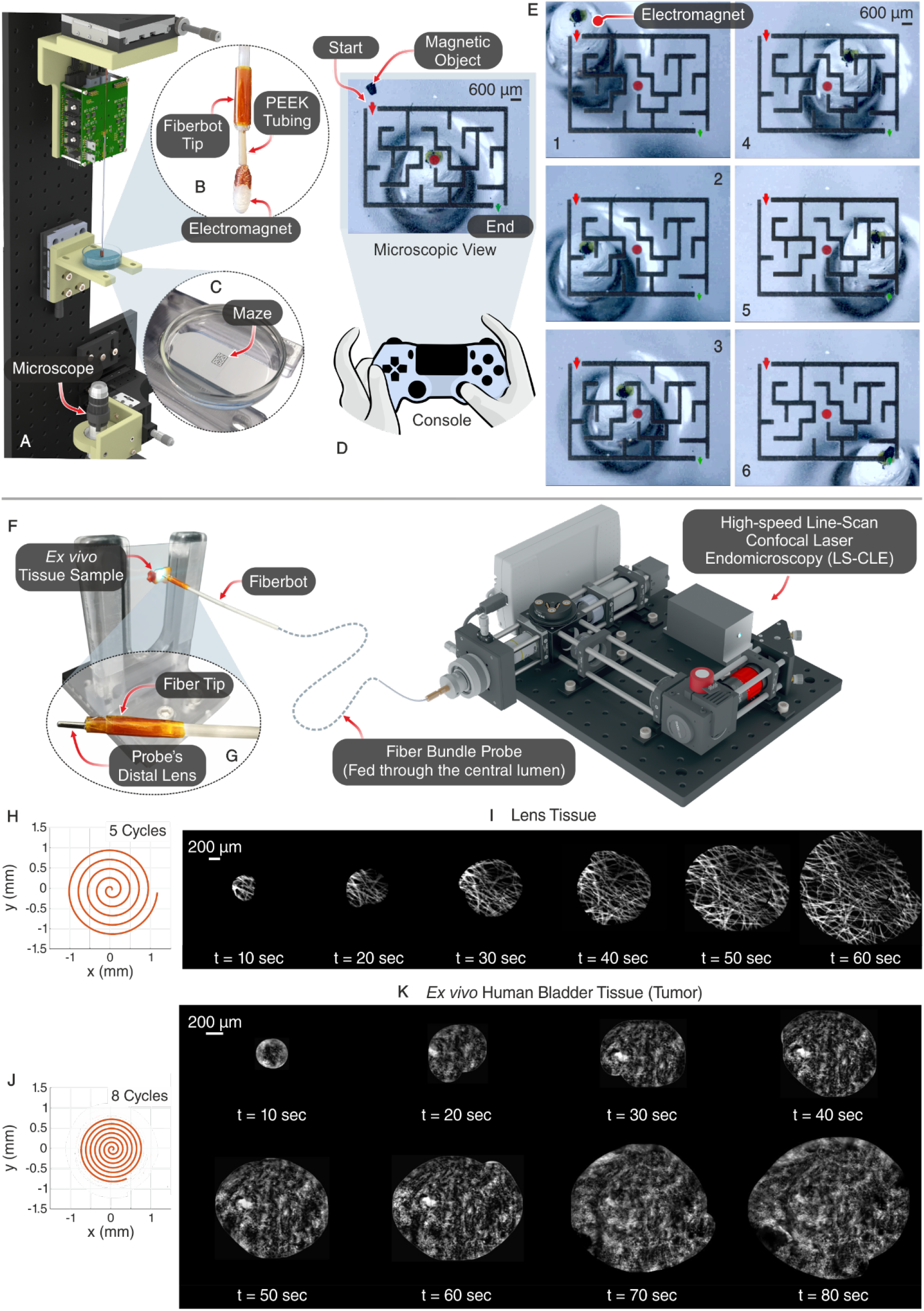
Magnetic tele-micromanipulation system and integration with the LS-CLE platform. (**A**) Vertically oriented experimental setup developed to micromanipulate magnetic objects. (**B**) Fiber coupled with an electromagnet and a multi-lumen PEEK tube to guide the electromagnet’s copper wires. (**C**) A water-filled Petri dish with a maze pattern printed on transparent paper attached to it. (**D**) Microscope view of the maze pattern and magnetic object as visual feedback to the user (top), alongside a joystick controller to teleoperate the fiber (bottom). (**E**) A photographic time series of manipulating the magnetic particle through the maze. (**F**) Overview of the integration between the LS-CLE system and the electrothermal fiber via a fiber bundle probe. (**G**) Close-up image of the thermally drawn fiber embedded with a probe through its central channel. (**H, J**) Desired spiral path of 5 and 8 cycles to be followed by the fiber. (**I, K**) Growing mosaics reconstructed at every 10 s interval from the acquired images while scanning a lens tissue and an *ex vivo* human urothelial carcinoma tissue respectively.

### Integration with the LS-CLE platform

The performance and control of the fiberbots also suggest many possible applications in surgery and medicine. Figure 4 (F to K) shows an exemplary application where we employed the fiberbots to precisely maneuver an imaging fiber bundle, a flexible tool of a microscopic imaging system used by surgeons and pathologists to intraoperatively detect and distinguish cancer and normal cells at surgical margins *(38, 39)*. This integration addresses the difficulty in assessing tissue morphology using the fiber bundle’s small field-of-view (325 μm in this case) by moving it over an area of several square millimeters while individual frames are stitched together to form a mosaic in real-time *(40)*. To implement this, we inserted the fiber-bundle (0.9 mm outer diameter, 3.5 μm resolution) through the fiberbot’s central channel (Fig. 4G) and optically coupled its proximal end to a portable and compact line-scan confocal laser endomicroscopy (LS-CLE) system. The probe was then moved at its distal end along a preoptimized spiral path at a scan time of 40 s (lens tissue) and 140 s (*ex vivo* tissue from the urinary tract surface) (movie S6). Based on the growing mosaics reconstructed at every 10 s from the acquired images, it can be seen that the integrated device was able to yield contiguous mosaics with high image quality (Fig. 4, I and K). In the lens tissue case, the hyper-fluorescent fiber structures were visible throughout the scan area (Fig. 4I). Similarly, for the urothelial carcinoma tissue, despite it being a soft surface, the large-area mosaic was reconstructed without any gaps showing features like the hyperfluorescent clusters of the nuclei with no defined organization and fibrovascular stalks with a distorted vasculature (Fig. 4K). These preliminary results demonstrate the feasibility of obtaining histology-like images without removing or destroying any tissue structure.

### Integration with the REIMS platform

Our next demonstration focused on presenting the fiberbots’ potential to combine several components to perform a surgical intervention with cellular-level precision. We coupled the fiberbot with a therapeutic tool (a surgical CO_2_ laser source) and a diagnostic tool (Rapid Evaporative Ionization Mass Spectroscopy (REIMS)) to showcase precise tissue ablation and identification of a 10 μm thick mouse brain section *(41)*. REIMS is a mass spectrometry technique originally developed to provide rapid and sample preparation free, molecular histology-based *in vivo* tissue diagnostics. The technique commonly utilizes electrosurgical devices to ablate tissues, with the resulting plume collected by a mass spectrometer to analyze its chemical composition. The REIMS platform was previously employed in identifying bacterial strains *(42, 43)*, determining food authenticity *(44, 45)*, and performing fundamental research on cell lines for pharmaceutical purposes *(46, 47)*. The working principle of the REIMS technique is described as follows: a given aqueous sample (e.g., tissue samples, bacteria, cell lines, plants, food samples) is exposed to rapid heating of its water content. This heating can be performed using Joule heating (e.g., electrosurgical devices), ultrasonic, laser, or contact heating. When the water content of the tissues is rapidly heated up and evaporated, the biological material is ejected as an aerosol rich in chemical information representative of the original samples. This aerosol is then directed towards a mass spectrometer for chemical analysis.

First, we introduced the previously used flexible PBG laser fiber through the fiberbot’s working channel to ablate the *ex vivo* tissue sample and attached a silicone suction tube to the outer surface of the fiberbot to simultaneously collect the generated aerosol for analysis by the REIMS system (Fig. 5B). The tissue sample was ablated by autonomously moving the embedded laser fiber along a preoptimized path of parallel lines at an average velocity of 0.7 mm/s and scan time per line of 6.3 s, during which data was collected through the suction tube at a rate of 5 scans per second. Figures 5D and 5E demonstrate the mass spectra obtained from the tissue regions of the dentate gyrus and fiber tracts, respectively, showing a comprehensive molecular coverage of small molecules, fatty acids, and structural phospholipids. In addition, for better visualization of the medical imaging capabilities of the integrated system, we generated a heat map image of the PA 36:2 [M-H]-phosphatidyl acid structure lipid ion from the data acquired using the REIMS system, which was then aligned and overlaid onto the optical image of the tissue sample (Fig. 5, F to J). The green-yellow bars represent a higher abundance, and the purple regions hint at the absence of the molecule. The abundance of the PA 36:2 [M-H]-ion shows a corresponding overlap with the fiber tracts region of the mouse brain, which correlates well with the *a posteriori* histological examination, in opposition to the low-intensity areas (dentate gyrus). The fiber tracts are white matter regions composed of myelinated axons, while the dentate gyrus is composed of grey matter made up of cell bodies. This explains the difference in mass spectra between these two regions as the biochemical composition varies in both ion species and relative intensities. These results provide a strong basis for developing the proposed system into a clinical tool combining robotics, diagnostics, and therapeutics, forming the core of an autonomous surgical system that could perform accurate, automated, and molecular histology-guided surgical resections.

**Fig. 5.**
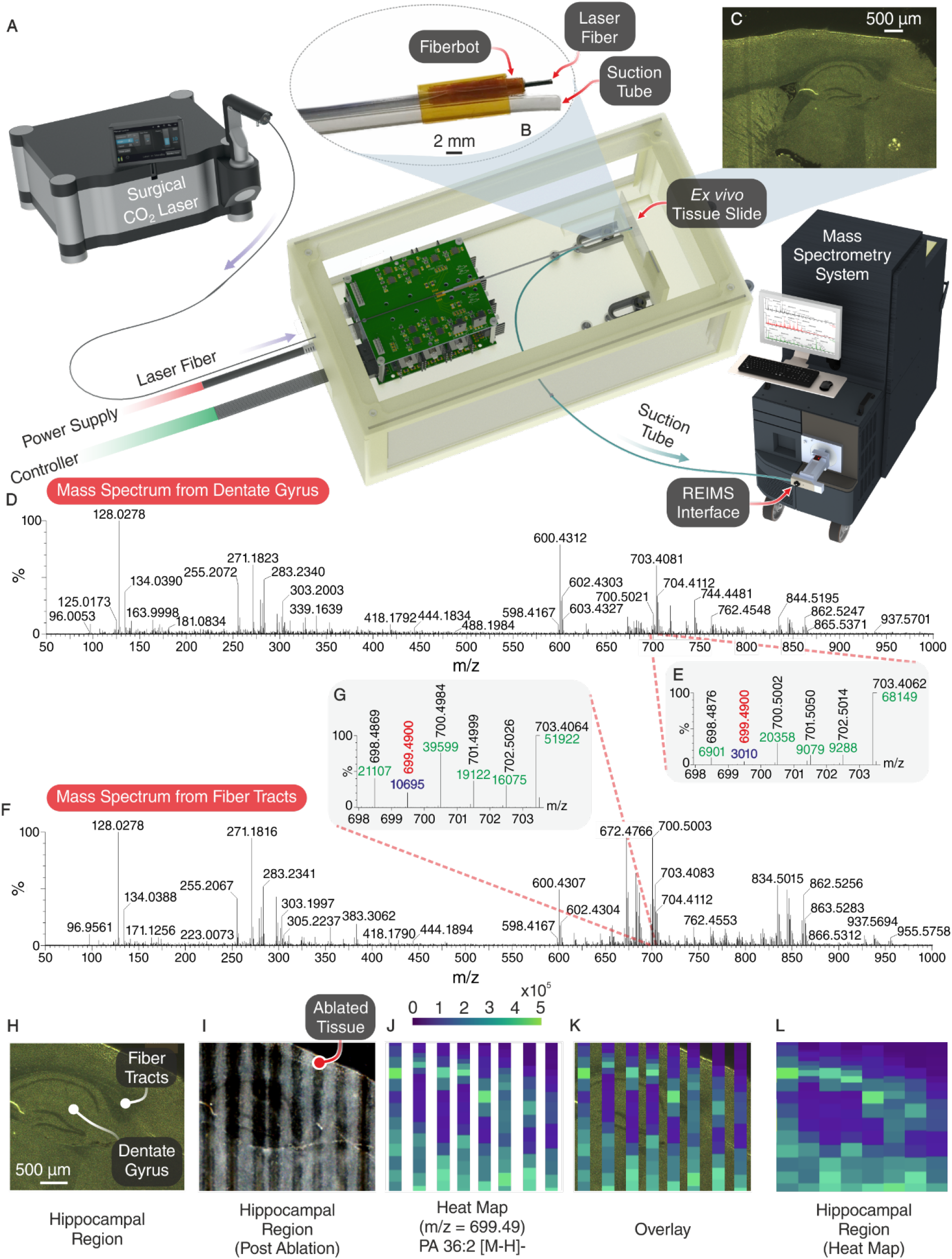
Integration with the REIMS platform. (**A**) Overview of the integration between the surgical CO_2_ laser, electrothermally actuated fiber, and REIMS interface. (**B**) Close-up image of the thermally drawn fiber embedded with a laser fiber through its central channel and an externally attached suction tube. (**C**) Optical image of a mouse brain’s sagittal section of the hippocampal region. (**D, F**) Mass spectra corresponding to the dentate gyrus and fiber tracts which shows a difference in the ion species and intensities corresponding to the grey matter distribution in the former and white matter in the latter. (**E, G**) Close-up of the mass-to-charge ratio (m/z) 699.4900. (**H**) Cropped optical image of the hippocampal region. (**I**) Optical image following the laser ablation of the tissue sample (parallel ablated lines created by moving the laser fiber using the electrothermally actuated fiber). (**J**) Single ion image heatmap for m/z 699.4900 tentatively assigned as PA 36:2 [M-H]-phosphatidyl acid, showing abundance in the fiber tract and relative absence in the dentate gyrus. (**K**) Overlay of heat map onto the optical image of the hippocampal region. (**L**) Stitched heatmap.

### Selective actuation of longer fiberbots and Da Vinci® integration

In an attempt to reveal the true potential of high aspect ratio materials, we explored a different fiberbot design to extend the fiber’s navigational capabilities and range of applications while maintaining the previously demonstrated benefits of scalability, precision, and low-power operation (Fig. 6, fig. S9). Contrary to the previous fiberbot design, three triples of equidistantly spaced wires (copper, stainless-steel, copper) were co-fed and embedded longitudinally within a 1.2 m long fiber during fabrication. Employing three controllable sides instead of four sides simplified the overall design and control of the fiberbot. The selective actuation of the fiberbot was realized by intertwining the resistive stainless-steel wire with one of the conductive copper wires at the distal end, followed by exposing the same resistive wire and the other conductive wire at a selected point along the length of the fiber by scraping the polymeric encapsulation with a microtome blade. Subsequently, conductive silver paint was applied to both points to ensure reliable electrical contact (Fig. 6, A to F). We then placed the fiber device in a tortuous configuration with radii of curvatures ranging from 12 to 15 cm to exemplify its flexibility (Fig. 6, G and H), demonstrating the potential endoluminal deployment of the fiber device through a flexible scope. Movie S8 shows the tele-micromanipulation of the selectively actuated fiberbot to reach four different targets. Using the same principle, we fabricated a bespoke holder to integrate the fiberbot with the Da Vinci® robotic surgical system and scan along the contour of a simulated tumor tissue (movie S9). This preliminary experiment outlines the versatility of the fiberbot and its ability to work hand-in-hand with commercially available surgical platforms, thus enabling the integration of multifunctional devices and enhancing access capacity (Fig. 6, K to N).

**Fig. 6.**
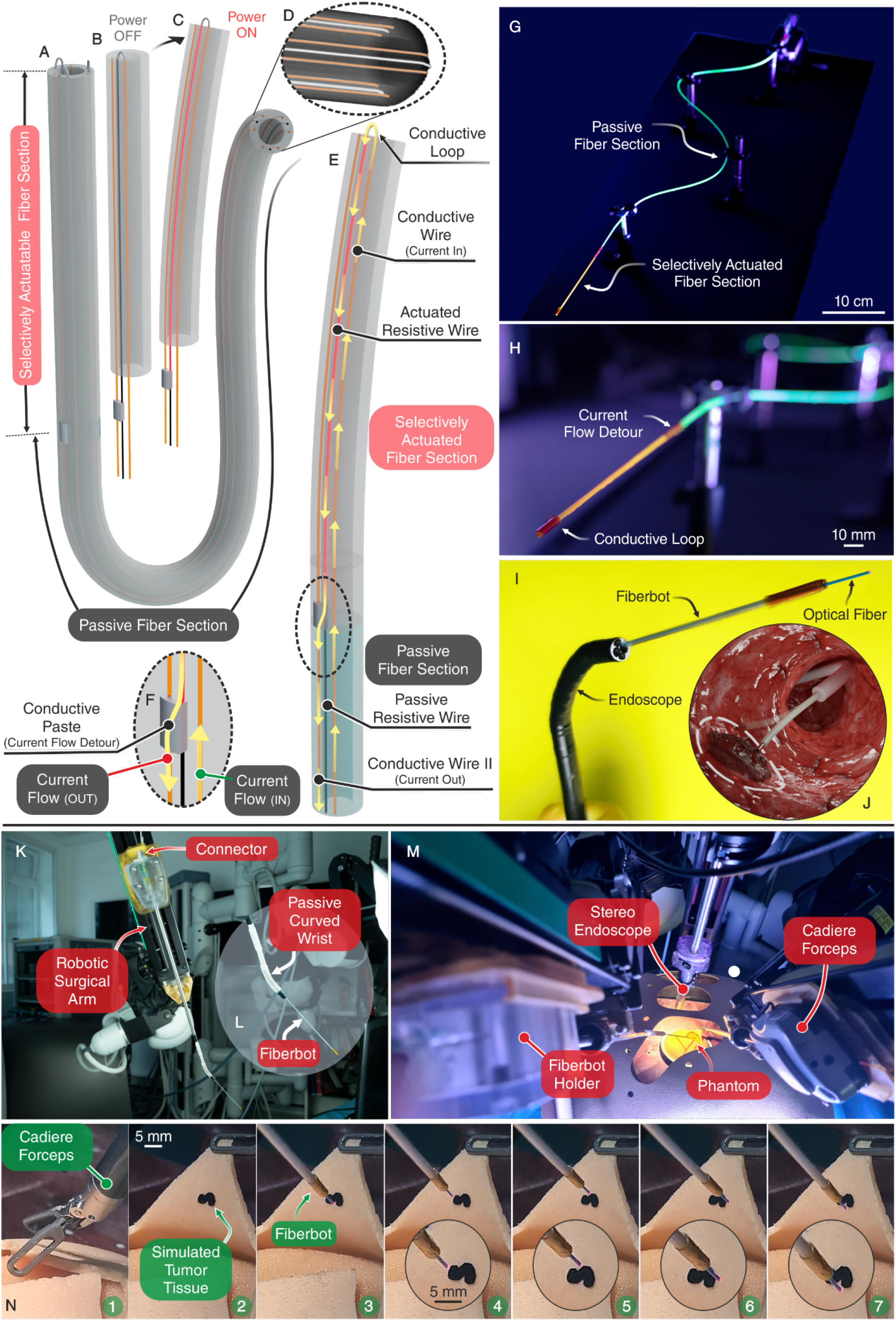
Selective actuation of longer fibers and integration with the Da Vinci® surgical robot. (**A**) Illustration of the long fiber with a selectively actuatable section and a passive section. (**B, C**) Illustration of the selectively actuatable section with power off (**B**) and power on (**C**). (**D**) Micro-computed tomography (μ-CT) scan of the 9-wire selectively actuated fiber. Wires highlighted in orange are (46 AWG) copper wires, and wires highlighted in silver are (44 AWG) stainless-steel wires. (**E, F**) Illustration of the wire connections at the distal end and selected proximal point, and the current flow path. (**G, H**) Images of the bent fiber with fluorescent spray paint painted over the selectively actuatable section (orange) and passive section (green). (**I**) Image of the selectively actuated fiber being inserted through the working channel of an endoscope, demonstrating flexibility and integrability. Similarly, an optical fiber is subsequently inserted through the central channel of the actuated fiber. (**J**) Conceptual image of the assembly presented in (**I**) in the removal of colon polyps. (**K, L, M**) The integration between the Da Vinci® surgical robot and the fiber actuator with a fiber holder and a passive curved wrist. (**N**) A photographic time series of scanning along the contour of a simulated tumor tissue with an embedded optical fiber.

### In vivo validation with a porcine model

The series of demonstrations presented above showcased our fiberbot’s capabilities in providing high-precision manipulation, well-controlled ablation, and enhanced tissue analysis. Although promising, evaluating the fiberbot’s performance in *in vitro* and *ex vivo* settings does not fully characterize and gauge the device’s clinical efficacy and safety outcomes. Therefore, to demonstrate that the precision of the fiberbot can be delivered safely under realistic *in vivo* conditions, we conducted animal testing using a porcine model (Fig. 7).

**Fig. 7.**
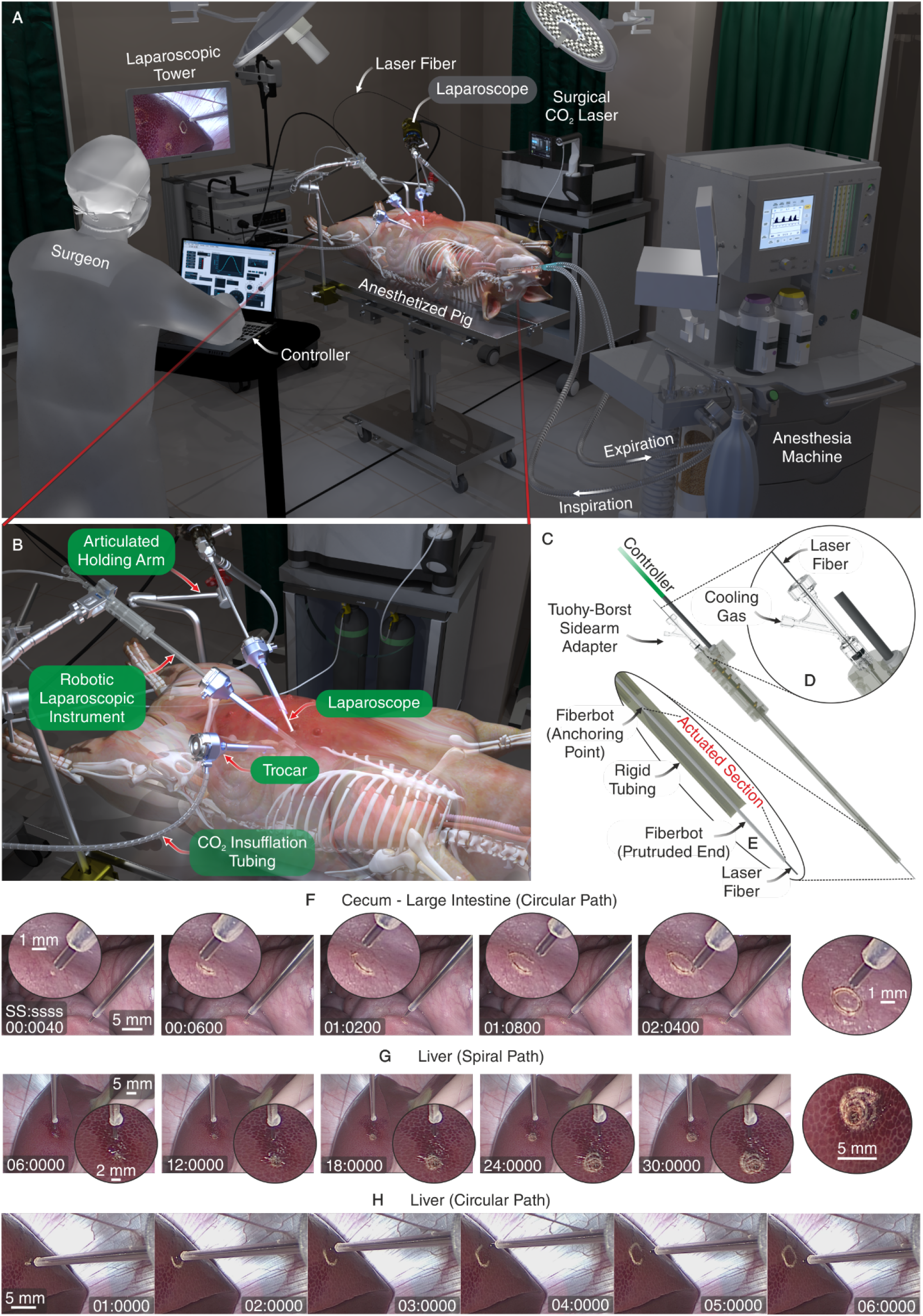
*In vivo* demonstration of the laparoscopic robotic instrument in a porcine model. (**A**) Graphical overview of the experimental setup. The setup features an endotracheally intubated pig put under general anesthesia in the supine position, an anesthesia machine employed to induce safe apnea phases (i.e. suppress respiratory-induced motion) during the robotically-controlled ablations, a surgical CO_2_ laser source which is optically coupled to the flexible PBG laser fiber, a laparoscopic tower which is optically coupled to the laparoscope, and a control unit which the surgeon controls to teleoperate the laparoscopic robotic instrument. (**B**) Close-up view of the CO_2_ insufflated abdominal cavity, inserted trocars, and the subsequently advanced laparoscope and robotic instrument. The articulated arms are used to position, orient, and secure the laparoscope and robotic instrument in place. (**C**) Illustration of the robotic instrument’s handle which hosts the electronics interface and the rigid tubin acting as a conduit for the fibrebot. (**D**) Close-up view of the Tuohy-Borst adapter which facilitates the simultaneous insertion of the laser fiber and injection of cooling gas through the fiberbot’s central channel. (**E**) Cross-sectional close-up view of the actuated section of the fiberbot and the rigid tubing’s distal end. In this configuration, the anchoring point of the (11.5 cm long) actuatable section of the fiberbot is recessed 8 cm inside the rigid tubing. For the recessed section of the fiberbot to move freely without any constraints, the inner diameter of the rigid tubing’s distal end is enlarged from 2 mm to 9 mm. Recessing the actuatable section of the fiberbot facilitates the safe and controlled insertion of the instrument through the trocar, and ensures that the overall length of the instrument is within a similar footprint to standard laparoscopic instruments (i.e. 35 cm long). (**F, G, H**) A photographic time series (captured using the laparoscope) of the fiberbot’s simultaneous actuation and ablation (during the safe apnea phases) of a: (**F**) circular path in the cecum, (**G**) spiral path in the liver, (**H**) circular path in the liver. Note: The scales were estimated using the closest point of the laser fiber to the tissue.

In line with our previous demonstrations, we developed a laparoscopic robotic instrument using our fiberbot to safely deliver precise tissue ablations during single-port, multi-quadrant surgeries that would typically be employed in the abdomen, thorax and head and neck. In this exemplar, the robot was used to access the liver and intestine within the abdomen. The standalone instrument comprises an electronic interface within the handle portion, onto which the embedded wires are soldered and anchored. Attached to the instrument’s handle is a 35 cm long rigid tube that acts as a conduit for the fiberbot which protrudes 35 mm out of the tube (Fig. 7, C and E). At the proximal end of the handle lies a Luer adapter that connects a Tuohy-Borst side-arm adapter to the fiberbot (Fig. 7D). The Tuohy-Borst adapter facilitates the simultaneous: (a) insertion of the flexible PBG laser fiber through the fiberbot’s working channel, (b) injection of helium gas around the laser fiber to cool down the fiberbot’s average temperature, and (c) prevention of the backflow of the injected helium gas. As discussed in the Supplementary Materials, the protruding end of the fiberbot was tested for electrical safety by immersing it in a conductivity standard solution (0.1413 S/m) which, on average, mimics the electrical conductivity of tissues in abdominal organs (fig. S10). It was found that no leakage currents flow from the fiberbot to the surrounding conductive solution.

For the *in vivo* experiment, the pig was endotracheally intubated and put under general anesthesia in the supine position, as shown in Fig. 7A. The trial commenced by inserting an 11 mm trocar into the abdomen through an incision made along the umbilicus. The bladeless obturator of the trocar was then removed, and CO_2_ gas (15 mmHg at a rate of 4 to 6 liters per minute) was insufflated into the abdominal cavity (pneumoperitoneum) through the trocar to create an adequate workspace between the viscera and abdominal wall. A laparoscope was then advanced through the trocar sleeve to visually guide the insertion of two additional 11 mm lateral trocars. Once all trocars were inserted, their bladeless obturators were removed, and the robotic instrument was introduced through one of the two trocar sleeves under the guidance of the laparoscope. Subsequently, both instruments (i.e., laparoscope and robotic instrument) were positioned, oriented, and secured in place using articulated holding arms (Fig. 7B). Note: In some cases, the laparoscope and robotic instrument are retracted and inserted through different trocars for better access and visualization of the target tissue.

Prior to actuating the fiberbot and performing ablations on the target tissue, we had to mitigate the inaccuracies introduced by the movement of abdominal organs caused by mechanical ventilation (i.e., respiratory-induced motion (RIM)). For example, during mechanical breathing, the liver’s RIM could range from 8 to 25 mm in one direction *(48 - 50)*. Therefore, if RIM is left unsuppressed, our instrument loses its accuracy and precision due to the time-varying offset between the fiberbot’s focus and the target tissue. Thus, making it challenging to locate and target exact lesions without risking damage to the surrounding healthy tissue *(51)*. Researchers over the years have developed machine learning, computer vision, and advanced control techniques to track and compensate for target tissue motion using teleoperated *(52, 53)* and autonomous robotic platforms *(54)*. Even though these approaches have proven their feasibility and effectiveness in the compensation of RIM, the focus of this study was to demonstrate the potential of the fiberbot as a standalone instrument without introducing additional equipment and complexity to standard laparoscopic surgeries. To achieve the objective above, we pursued a well-established and safe intraoperative technique called *“adequate preoxygenation with apneic oxygenation*.*”* This method utilizes the anesthesia machine to control respiratory movements during procedures that require higher precision (e.g., radiation therapy in breast cancer treatment *(55)*, coronary artery bypass grafting *(56, 57)*). In this technique, anesthesiologists start by preoxygenating (denitrogenating) the patient to 100% oxygen (O_2_) for a few minutes before switching off the mechanical ventilator and inducing a long and safe apnea phase (i.e., periods with no breathing and RIM in intubated patients). During this apnea phase, the patient’s O_2_ flow is maintained while the physician performs the desired treatment. Anesthesiologists preoxygenate the lungs to increase the O_2_ reserve within the alveoli, thus increasing the safe apnea time. Otherwise, the persistence of nitrogen and the accumulation of CO_2_ during apnea could speed up the onset of hypoxemia (i.e., lower levels of oxygen in the blood than normal). The safe apnea time, i.e., the duration until critical arterial desaturation occurs following the cessation of ventilation, can be up to 8 minutes in healthy adults, 5 minutes in moderately ill adults, and 2.7 minutes in obese adults *(58)*. Critical arterial desaturation occurs when O_2_ saturation (SaO_2_) falls below the upper inflection point on the oxygen-haemoglobin dissociation curve, beyond which a further decrease in the partial pressure of O_2_ (PaO_2_) results in a rapid decrease in SaO_2_. Note: SaO_2_ is the percentage of oxygen in the arterial blood, PaO_2_ is the oxygen pressure in the arterial blood, and the oxygen-haemoglobin dissociation curve is the curve that correlates oxygen saturation across a range of oxygen pressures. Moreover, the timeframe of minutes is adequate to execute the desired laparoscopic ablations using the fiberbot. Readers are encouraged to review Lyons *et al*.’s article, which summarizes the underlying mechanisms of gas exchange during apneic oxygenation and its various clinical applications *(59)*.

After identifying different target organs and tissues to ablate, our instrument’s efficacy was evaluated during the apnea phase by commanding the fiberbot (with the flexible PBG laser fiber introduced and activated, and helium gas injected at 50 Pound-force per square inch (psi)) to follow several pre-defined desired paths (e.g., circle and spiral). In addition, we monitored the pig’s physiological variables (i.e., SaO_2_, heart rate, temperature, inspiratory CO_2_, end-tidal CO_2,_ inspiratory O_2_, expiratory O_2_, inspiratory and expiratory inhalational anesthetic agents) during the experiments to ensure and demonstrate the safety of apneic oxygenation. Figures 7F and 7H show two photographic time series of the fiberbot following different circular paths while it simultaneously and safely ablates the cecum (part of the large intestine) and liver tissue, respectively. Figure 7G shows a similar series of the fiberbot’s simultaneous actuation and liver tissue ablation of a spiral path. For the cecum, the ablation of the circular path took approximately two seconds with no additional complications and changes in the pig’s physiological condition. Similarly, for the liver, the fiberbot took 30 seconds to complete the spiral path and 6 seconds to complete the relatively larger circular path; both with no adverse events (e.g., critical arterial desaturation and CO_2_ accumulation) observed during and after the experiments (Movie S10).

Under the same *in vivo* conditions, we demonstrated the thermal safety of the instrument by conducting a series of experiments where the fiberbot was actuated (i.e., one side of the tube is heated) and held against the *in vivo* tissue for a few seconds. During these experiments, we kept the: (a) flexible PBG laser fiber advanced through the central channel of the fiberbot, and (b) helium gas injected at a constant pressure of 50 psi to cool down the fiberbot. Using a thermocouple, we monitored and adjusted the outer surface temperature of the fiberbot by controlling the (electrical) input power. We tested four different temperatures (42±1°C 43±1°C, 55±1°C, and 60±1°C) and held the heated side of the fiberbot against liver and spleen tissue for two different exposure times (5 and 10 seconds) as shown in Figures 8A and 8B. The exposed tissues were subsequently excised for the histopathological assessment of thermal injury (Fig 8, C and D). The histopathological analysis showed no morphological signs of coagulation due to thermal injury in all tissue planes of the liver and spleen (Fig 8E). These results validate the efficacy and safety of the fiberbot in realistic *in vivo* conditions as a robotic laparoscopic instrument.

**Fig. 8.**
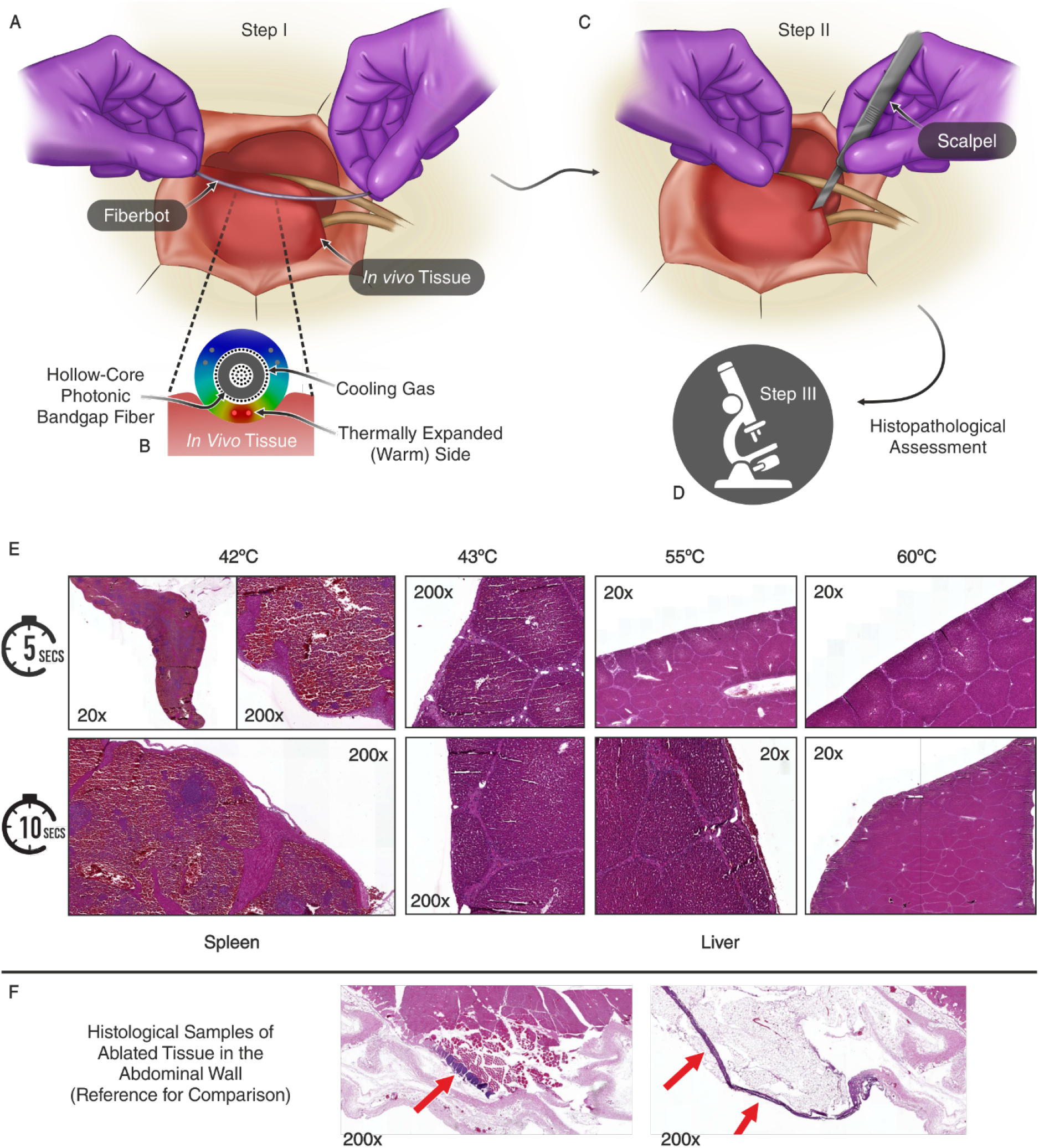
Histopathological assessment of thermal injury. (**A**) Graphical illustration of the first step in the experiment, where the actuated side of the fiberbot is held against the *in vivo* tissue for a few seconds. (**B**) Cross-sectional closeup view of the fiberbot and its interaction with the *in vivo* tissue. The fiberbot is embedded with the flexible PBG laser fiber and injected with a constant pressure of helium gas to cool it down. Note: The laser fiber employs a similar cooling technique using helium gas. (**C, D**) Graphical illustration of the second and third steps in the experiment, where the exposed tissue are excised for the histopathological assessment of thermal injury. (**E**) Optical images of eight 4 μm thick and hematoxylin and eosin (H&E) stained spleen and liver tissue slides. The acquired tissue samples were exposed to different temperatures (42°C, 43°C, 55°C, and 60°C) for differing durations (5 and 10 seconds). No morphological signs of coagulation due to thermal injury were observed at the capsular surface or parenchyma of the liver and spleen tissue samples. 20x and 200x indicate the magnification of the objective lens used to capture these images. (**F**) Optical images of two ablated tissue samples of the abdominal wall which demonstrate thermal injury of connective & muscle tissue at the surface of the peritoneum (i.e., serous membrane forming the lining of the abdominal cavity). These examples are presented as a reference for comparison with our experimental results.

## DISCUSSION

In this work, we presented a rapid prototyping approach that simplifies the production of hundreds of meters of inexpensive, bespoke, precise, and integrable actuators that can readily function. The fibers we produced at scale comprise flexible and compact polymer-metal composites that can move with high-precision by asymmetric thermal expansion. The flow of electric current through the longitudinally embedded metallic wires generates heat, which leads to a localized increase in the temperature of the wire and its enveloping polymer; and their subsequent thermal expansion. Unlike previous research efforts that utilize thermally responsive polymers to steer tubular structures (e.g., hydrogels that shrink *(60)* and liquid crystal elastomers (LCEs) that change in phase *(61)* when heated), we functionalized a class of thermoplastic polymers that are often: (i) not classified as thermo-responsive polymers and (ii) disregarded in the actuation of soft robots due to their relatively large flexural rigidity (i.e. the stiffness of the material when its subjected to bending) when compared to softer polymers. Moreover, we capitalized on their stiffness and intrinsic (positive) thermal expansion property to create stable actuators that can omni-directionally move with high-precision (over four orders of magnitude) and negligible hysteresis. Although PC and stainless-steel were employed in our study, the findings can be reproduced using thermally drawn fibers with different cross-sectional designs, made of similar thermoplastic polymers (e.g. polysulfone, polyetherimide, poly(methyl acrylate)) and metallic wires (e.g., nickel-chromium, nickel-titanium).

After characterizing and controlling the millimeter-scale fiberbots, we proposed and demonstrated several applications in surgery and medicine, fields that demand micron-level precision from high-aspect-ratio devices. The fiberbot’s tubular structure facilitated these applications by seamlessly integrating therapeutic and diagnostic platforms, thus providing high-precision manipulation, ablation, and enhanced tissue analysis *in situ* down to the cellular level. We also used the fiberbot to perform challenging micromanipulation tasks that may inspire new applications. For example, beyond the scope of the work presented, we envision that these fiberbots could potentially be used as flexible 3D printing nozzles for printing small objects in confined spaces *(62)*. Further demonstrations with extended fiberbot lengths highlight the possibility of using such actuators in challenging sites that demand flexibility and miniaturization.

Through the preclinical testing, we demonstrated the efficacy of the fiberbot as a minimally invasive robotic instrument in an *in vivo* setting. This demonstrated that the fiberbot has clinical utility and applications in tissue imaging, molecular sensing (REIMS) and the delivery of therapeutic lasers for clinical benefit. The advancements in endoscopic and microrobotic interventions are being driven by the requirement for the early diagnosis and precision therapeutics of cancers of poor prognosis (e.g., brain, ovarian, lung, hepatobiliary and pancreas) and cardiovascular diseases with high operative morbidity. Moreover, the presented millimeter-scale robotic systems have the potential to improve the safety and reliability of these interventions.

Our histopathological assessment showed that although the fiber is electrothermally actuated and can reach high outer surface temperatures (up to 60°C with helium gas injected), it does not cause any thermal injury to the surrounding tissue for two different exposure times (i.e. 5 and 10 seconds). This is primarily due to the low thermal conductivity of the fiber’s encapsulating polymer (PC: 0.22 W/(m·K)), which acts as a buffer for heat transfer and limits the amount of heat transferred to the surrounding tissue. Conversely, if the same set of experiments were conducted using a metallic instrument (e.g., Platinum: 71.6 W/(m.K)), which had the same outer surface temperature, the possibility of it causing a thermal injury would have increased. Moreover, our electrical safety experiments corroborate that the fiberbot is safe and reliable for *in vivo* use. Therefore, we envisage that the proposed instrument can give surgeons greater control in minimally invasive laparoscopic surgeries.

However, there are some potential technical challenges to consider for further clinical translation of the fiberbot. To begin with, the fibrebot’s length affects its response to thermal expansion, i.e., the shorter it is, the smaller its range of motion and reachable workspace. Therefore, in clinical applications where the target tissue is in confined and tortuous spaces, the relatively long actuatable section of the fiberbot would impede it from being positioned normal to the target tissue. In addition, although the embedded straight metallic wires expand, their thermal expansion is relatively smaller when compared to the enveloping PC, which also restricts the overall motion of the fiberbot.

One possible way to achieve a larger range of motion for a shorter fiberbot length would be to select thermoplastics with similar mechanical properties and thermal conductivity but with higher thermal expansion coefficients. Other approaches include adding additional channels within the thermally drawn fiber to locally cool down the passive side of the fiberbot and increase the temperature difference between the actuated and passive sides. By cooling down the passive side, we limit its thermal expansion and increase the effective thermal expansion of the actuated side and, consequently, the overall bending of the fiberbot. To overcome the restriction caused by the embedded metallic straight wires, substituting them with wave-shaped wires (such as those used in cochlear implants) or coiled wires could potentially increase the fiberbot’s range of motion.

## MATERIALS AND METHODS

### Preform fabrication

The preform fabrication process commenced by defining the geometry of the preform using 3D Computer-Aided Design (CAD) software (SolidWorks 2020, Dassault Systèmes, France). The preform design was then 3D printed by feeding PC (Ultimaker PC Transparent, 2.85 mm, Netherlands) filaments into a commercially available fused deposition modeling (FDM) printer (Ultimaker 3 Extended, Netherlands). The warping of the 3D printed preform was prevented by: (a) using an adhesive layer (DIMAFIX, 3D GBIRE, Lancashire, United Kingdom), which was sprayed onto the glass build plate, and (b) fitting the printer with an upper enclosure and a door (Accante cover Ultimaker 3 Extended, 3D GBIRE, Lancashire, United Kingdom) for a constant printing chamber temperature during the printing process. The dimensions of the 3D printed preform were 40 mm in diameter and 200 mm in length. The central channel was 24 mm in diameter, while the smaller side channels were 2 mm in diameter. For the 3D printing parameters, refer to Table 1 of reference *(63)*.

### Fiber drawing

The preform was coupled to a PEEK 30% glass fiber reinforced preform holder (Ketron Peek Glass Fiber Reinforced Rod, 20 mm in diameter, Bay Plastics Ltd, UK) and a wire feeding platform comprising eight bobbins of 50 μm bare stainless-steel wires (44 AWG (0.05 mm), The Crazy Wire Company, UK). Since the preforms are printed layer-by-layer and have lower tensile strengths than molded or extruded solid structures, their desired viscosity, when drawn, is expected to be lower than that of solid preforms (*64*). By setting a high initial draw temperature of 265°C, the preform’s viscosity dropped several orders of magnitude (fig. S11B) and necked down under (a) the constant tension of 55 grams and (b) the weight of the lower end of the preform, which formed the neck-down region. Note: The actual temperatures inside the furnace (TV-05 Solid Tubular Furnace, The Mellen Company, USA) are lower, refer to (*65*) for the furnace calibration. When the lower end of the preform exited the furnace, it was attached to a pulling system, and the draw temperature was lowered gradually to 220°C. In the selectively actuated fiber case, the wire feeding platform was extended to accommodate nine bobbins, where six bobbins of 40 μm bare copper wires (CU005200, 46 AWG (0.04 mm), GoodFellow, UK) and three bobbins of 50 μm bare stainless-steel wires were fed simultaneously.

### Experimental setup for fiberbot actuation characterization

The enclosed setup in fig. S12A was built to provide a stable characterization environment for the proposed fiberbot actuation mechanism with minimal external interference. An XYZ stage (LT3/M-XYZ translation stage, Thorlabs, USA) was used to adjust the fiber actuation PCB assembly’s position to ensure that the fiberbot’s distal tip is aligned with an optical triangulation laser sensor (LK-G5000, Keyence, Japan). This sensor was employed to accurately measure the displacement of the fiberbot tip in a specific direction. With the characterization setup completely enclosed, the proposed fiberbot system’s resolution, accuracy, and repeatability were thoroughly assessed. For Fig. 2C, the relationship between input electric power and fiberbot tip displacement was characterized by performing an input power sweep from 0 W to 0.95 W in approximately 95 mW incremental steps for the fiber with an outer diameter of 1.65 mm and 12 cm in length; and from 0 W to 1.12 W with approximately 160 mW increments for the fiber with an outer diameter of 2.00 mm and 10 cm in length. The input power was maintained for each measurement for 2.5 minutes, and the average tip displacement of the last 1.5 minutes from the 2.5 minutes period was calculated. For the (2.00 mm and 10 cm) fiberbot, the same experiment was repeated for five times, and the mean and standard deviation of each set of measurements for the same input power were plotted. In Fig. 2D, using the (2.00 mm and 10 cm) fiberbot, the positional resolution was measured by also performing an input power sweep with ≈ 13 mW increments, intermitted by 2.5 minutes. For the contact force measurements shown in Fig. 2E, the (2.00 mm and 10 cm) fiberbot tip was placed against a flat-tipped force sensor (FT-S1000000 μN, FemtoTools, Switzerland). The sensor was mounted on the top of a 25 mm XYZ translation stage (PT3/M, Thorlabs, USA) and in close proximity to the fiberbot setup. To accurately align the sensor with respect to the fiberbot, a digital microscope (VHX-6000, Keyence, Japan) was employed, as shown in Fig. 2A. Similar to the previous characterizations, the force measurements (at zero displacements) were acquired by performing a power sweep from 0.105 W to 1.155 W with ≈ 105 mW increments. The input power was also maintained for each measurement for 1 minute, and the average contact force for the last 30 seconds of the single minute period was calculated. The same experiment was repeated five times, and the mean force and its standard deviation for each input power were plotted.

Additional characterization experiments were performed to further assess the performance of the proposed system by making use of additional structures/devices and setup re-arrangements. One such experiment consisted of exploring the relationship between the length of the actuated fiberbot and its corresponding tip displacement. A clamp was employed to tighten the fiberbot at different points, thereby ensuring that only part of the fiberbot could move freely during actuation. The same triangulation laser sensor was used to measure the displacement of the fiberbot tip in the horizontal direction. As demonstrated in Fig. 2G, a power sweep from 0 W to 1.449 W with ≈ 161 mW increments and intermitted by approximately 2.5 minutes was performed to capture the effect of the free length of the fiberbot on its tip displacement. The thermal images shown in Figs. 1H, 2H, and 2I were captured using a highly sensitive medium-wave infrared (MWIR) camera (FLIR X6900sc MWIR, Teledyne FLIR LLC, USA) to: (a) visualize the temperature distribution profile along the fiberbot’s length (Fig. 1H), (b) visualize the effect of different fiberbot lengths on its displacement (Fig. 2H), and (c) compute the temperature gradient between the opposing sides of the fiberbot (actuated side and passive side) as a function of the applied power for horizontal displacements (Fig. 2I). For Fig. 2I, the power sweep was performed in both directions from 0 W to 1.440 W with ≈ 180 mW increments.

Using the power measurement circuit (described in the Supplementary Materials, fig. S4), we characterized the change in resistance of the embedded wires due to fiberbot actuation by applying 10 V input steps, as shown in Fig. 2J. The displacement of the fiberbot tip was also acquired using the same laser displacement sensor. The velocity profiles presented in Fig. 2K were obtained by: (a) applying different voltage inputs, (b) measuring the displacements using the displacement sensor for each input voltage, (c) polynomial curve fitting of the measured displacement, (d) calculating the derivative of the curve fitted displacement with respect to time to get the approximate velocity profile of the fiberbot. Last, the hysteresis plot in Fig. 2L was obtained by performing an input power sweep from 0 W to 1 W and vice versa in 125 mW incremental steps.

### Open and closed-loop control mechanisms for fiberbot actuation

To implement different control strategies, the fiberbot was embedded with a hollow-core photonic bandgap (PBG) fiber (0.9 mm outer diameter, fiber was stripped of its polymeric jacket, OmniGuide, USA) to facilitate the tracking of its tip. The PBG fiber was coupled to a 650 nm light-emitting diode (696-1838, RS, UK) to track its tip using a digital microscope (VHX-6000, Keyence, Japan). Note: the visible light is guided by the polymer cladding of the PBG fiber, not the hollow core designed to guide the 10.6-micron wavelength. Furthermore, the bright polymer cladding of the PBG fiber tip was used to track the motion of the fiberbot tip in two dimensions (2D). An image grabber (AV.io HD™, Epiphan Video, Canada) was utilized to grab the images from the microscope’s display monitor into a portable workstation where a LabVIEW-based image tracking software and control system was implemented. A local descriptor for the PBG fiber tip was initially created in LabVIEW by using a low discrepancy sampling algorithm. This description data was stored in the system and subsequently used to search for the PBG fiber tip in each frame during the video acquisition. The IMAQ Match Pattern algorithm of the NI Vision 2018 library (version 2018, National Instruments, USA) was employed for this task. This algorithm provides the position of the steered PBG fiber in ***P***_***PBG_TIP***_ *= [X*_*PBG_TIP*_ *and Y*_*PBG_TIP*_] in each frame. After the first position is calculated, a region of interest around the PBG fiber tip is automatically generated (centered in ***P***_***PBG_TIP***_) to reduce the processing time and overall accuracy and reliability of the tracker *(66)*.

In the initial tests, an open-loop control mechanism was implemented to actuate the fiberbot to follow a specific path. By utilizing the measured power versus displacement curves, it was assumed that the power applied through a pair of stainless-steel wires is proportional to the displacement of the fiberbot tip as given below:

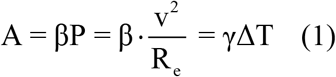

where A represents the amplitude displacement, P is the applied power, β is a constant derived from the power-displacement calibration, ΔT is the temperature difference between the opposing sides of the fiberbot, ϒ is a constant derived from the temperature-displacement calibration. The position of the fibrebot tip is represented by a 2D coordinate system, with the origin defined as the equilibrium position of the fiberbot (prior to actuation). Different paths (raster, square, star, spiral, and circle) were generated in Matlab (version 2020, Mathworks Inc., USA) as an ordered set of *[X and Y]* input coordinates to the open-loop controller. The flowchart shown in fig. S6 describes the methodology behind the open-loop controller developed in LabVIEW. Using the set of input coordinates, the first task of the controller is to find the region for the initial *[X and Y]* coordinates from a set of 8 regions spanning the 2D coordinate system (Fig. S6B). In vertical and horizontal displacements, only one pair of wires is powered based on the voltage calculated by equation 1. For diagonal displacements, two pairs of wires are powered based on the voltage ratio between the pairs controlling the direction of inclination. Following the first point, the remaining points of the path are executed one-by-one utilizing the aforementioned approach to determine the next pair(s) of wire(s) to be driven and the corresponding voltage(s). However, the results obtained by this open-loop controller have shown significant deviations from the ideal path (reference) due to the inherent slow thermal response of electrothermal actuation, as well as non-linearities such as the effect of gravity affecting the horizontal and vertical motions differentially. In order to compensate for these positional errors, the input paths were manually and iteratively optimized, as described in the main text and shown in figs. S6D to S6K.

For the closed-loop control strategies illustrated in fig. S7, four proportional-integrated-derivative PID VIs (version 2018, National Instruments, USA) were employed to simultaneously control the four pairs of wires. Herein, we used the opposing pairs of wires to control one dimension of the overall two-dimensional motion. For example: to move the fiberbot tip to the left-up direction, the input voltages across the left and top pairs of wires are decreased; concurrently, the input voltages across the right and bottom pairs of wires are increased. In the case of optical-based feedback, the fiberbot tip’s tracked position (process variable) was deducted from the input position (desired setpoint), and thus the deviation (error value) was calculated. Based on the calculated deviation, the manually tuned-PID controller and transfer function of the fiberbot (mapping displacement to voltage, equation 1) determine the voltage input for the pair of wires. On the other hand, for temperature-based feedback, we mapped the desired fiberbot tip displacement to the corresponding temperature difference between the opposing sides of the fiberbot using also equation 1 Then, the temperature difference measured (process variable) using the thermocouples (406-590, TC Direct, UK) was deducted from the input temperature difference (desired setpoint) to calculate the deviation (error value). Using the computed deviation, the manually tuned-PID controller and transfer function of the fiberbot (mapping temperature difference to voltage, equation 1) calculate the voltage input for the pair of wires. Moreover, four low-pass filters were also used to smoothen out the temperature signals detected by the thermocouples before feeding the measurements back to the controller (Fig. S7B).

Selective Actuation: Unlike the open-loop controller implemented for the previously described eight wires configuration, three equidistantly placed (120° apart) wires were used to steer the selectively actuated fiberbot. The motion of the selectively actuated fiberbot was controlled by dividing the 2D plane into three areas, as shown in Fig. S9E. As in the case of four pairs of wires, the first step was to find the relevant area of the desired point. Based on the defined area, the control system powers the relevant wire(s) for actuation. The contribution of each wire to the desired displacement was calculated using the sine law. For example: as shown in Fig. S9E, the green dot is the desired 2D position, located in area 1. The magnitude of the desired displacement vector is defined as A, whereas the magnitude of A’s projection along one of the actuatable axes is defined as B; similarly, the magnitude of A’s projection along the other actuatable axis (within the same area) is defined as C. In addition, the angle between C and A is defined as ∠ϕ. Moreover, the magnitudes of B and C can be calculated using the equation below:

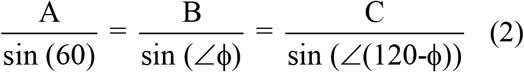

Based on equation 1, the desired power(s) and the voltage(s) are calculated.

### Integration with the REIMS platform

A Xevo G2-XS Quadrupole Time-of-Flight (Waters, MA, USA) equipped with a prototype REIMS source was used for the experiments. A CO_2_ laser source (OmniGuide, USA) was coupled with a flexible hollow-core photonic bandgap (PBG) fiber (0.9 mm outer diameter, fiber was stripped of its polymeric jacket, OmniGuide, USA) to guide CO_2_ (wavelength = 10.6 μm). The PBG fiber was subsequently inserted into the central channel of the fiberbot to perform simultaneous scanning and ablation of *ex vivo* tissue samples. The ensemble of these different components leads to the realization of an intelligent knife (iKnife) system for intra-operative diagnostic mapping *(67, 68)*. Furthermore, the fiberbot was placed in a bespoke enclosure to create a well-controlled and stable environment. The enclosure was designed using a 3D CAD software (SolidWorks, Dassault Systèmes, France) to accommodate the fiberbot, its electronic circuitry, and an *ex vivo* tissue slide holder. The enclosure comprised a 3D printed shell, printed in VeroClear material (standard quality, glossy finish) using a PolyJet printer (Objet 500 Connex, Stratasys Ltd, USA), and laser-cut 2 mm clear acrylic sheets (824-496, RS, UK), cut using a CO_2_ laser cutter (VLS 6.60, Universal Laser System, USA) to create a transparent enclosure. The adjustable tissue slide holder was also 3D printed in VeroClear. The holder was designed to be moved and mounted onto the shell’s base at different positions (i.e., adjustable) to accommodate different lengths of fibers. The tissue samples were ablated by moving the embedded laser fiber along a preoptimized path of parallel lines. A LabVIEW Visual Instrument (VI) (version 2018, National Instruments, USA) was developed to: (a) actuate the fiberbot, (b) switch on and off the laser source for ablation (“on” for the parallel lines part of the path, “off” for the intermediate motions between the lines), and (c) trigger the mass spectrometer to start the analysis of the aerosol. The mass spectral data were acquired between 50 – 1200 m/z in the negative ion mode at the rate of 5 scans *per* second.

### Mass spectral data processing and heatmap generation

The mass spectral data were acquired using Masslynx (version 4.1, Waters, USA) and were processed using Abstract Model Builder (version 1.0.2059.0., Waters Research Center, Hungary). First, the data were processed using data binning between 50 – 1200 m/z, with a bin width of 0.1 m/z. Subsequently, the data matrix was generated and exported from the software. The data were then imported into a custom-made script, where the mass spectral data acquired while ablating the tissue (i.e., laser on) were selected to generate the parallel line’s heat map. The time intervals as to when the laser was activated and deactivated were obtained from a data file generated by a dedicated LabVIEW VI that keeps a time-stamped log of the main control parameters. Next, the selected data were plotted in a two-dimensional heat map with equal pixel sizes for the 200 highest mass bins. Having equally sized pixels introduces inaccuracies to the heat map generated due to the change in velocity of the fiberbot as it moves along the parallel line path. This change in velocity (i.e., acceleration at the beginning and deceleration at the end of each line) stems from the relatively slow thermal response of electrothermal actuation. Since the mass spectral data is acquired at a fixed sample rate (5 scans per second) and the distance traveled during the acquisition of each scan is variable, there is a need to correct the size of each pixel based on the actual distances traveled by the fiberbot. These distances were obtained by processing the images of a video recording of the fiberbot in motion using MATLAB (version 2021, MathWorks, USA). A more accurate heat map was then generated by adjusting the pixel sizes based on the corrected distances at each mass spectral sampling position. The resulting pixel images were overlaid onto the optical image of the post-ablated *ex vivo* tissue slide.

### Description of the developed LS-CLE system

Figure S13 shows the schematics of the line-scan confocal laser endomicroscopy (LS-CLE) system. A full description of the system can be found in (*69*). The illumination consists of a compact laser diode delivering light at 488 nm (Vortran Stradus® 488-50, Vortran Laser Technology, USA) with a maximum output power of 50 mW. The output beam is expanded to about 4 mm by a 2X beam expander (GBE02-A, Thorlabs, USA). This is reflected off a galvanometer scanning mirror (GVS001, Thorlabs, USA) and enters a telescope consisting of a plano-convex cylindrical lens (LJ1695RM, f = 50 mm, Thorlabs, USA) and an achromatic doublet tube lens (AC254-050-A-ML, f = 50 mm, Thorlabs, USA). The cylindrical lens shapes the laser beam into a line of illumination, and a dichroic mirror (MD498, Thorlabs, USA) reflects this beam toward the 10X plan infinity-corrected microscope objective (RMS10X, Thorlabs, USA), which focuses it to a line across the proximal end of the fiber-bundle. While the developed fluorescence imaging system can be used with any fiber-bundle, for this work, a high-resolution flexible AQ-Flex19 fiber-bundle probe from Cellvizio (0.91 mm diameter, 325 μm field-of-view, 3.5 μm resolution, Mauna Kea Technologies, France) was utilized. The spatial resolution of 3.5 μm is sufficient to resolve normal epithelial cell nuclei that range in diameter from 5 to 10 μm. The working distance of the probes is about 60 μm making it suitable to image superficial cellular layers of the tissue under examination. A sterile plastic sheath with ∼50 μm thickness was kept between the probe tip and tissue surface to avoid contamination.

A standard fiber chuck (HFC005, Thorlabs, USA) and a fiber rotation mount (HFR001, Thorlabs, USA) were used to mount the proximal end of the fiber-bundle in the lens mount. The fiber-bundle probe transfers the laser light to the tissue with some *pixelation* due to the fiber core pattern. During scanning, a time-averaged power of ∼1.6 mW is delivered to the tip of the probe. Fluorescence emission from the tissue under examination is collected by the same fiber-bundle and transmitted by the dichroic mirror, fluorescence emission filter (FEL0500, Thorlabs, USA), and notch filter (NF488-15, Thorlabs, USA) to remove reflected 488 nm light. The proximal face of the bundle is de-scanned and imaged onto a rolling-shutter CMOS camera (FL3-U3-13S2M-CS, Point Grey Research (acquired by FLIR), Canada) using an air-spaced achromatic doublet (ACA254-075-A, f = 75 mm, Thorlabs, USA). The rolling shutter of the CMOS camera operates as a virtual detector slit that rejects most of the out-of-focus light, leading to optical sectioning. The camera is operated in free-run mode, allowing images to be acquired at the full frame rate of 120 fps by generating a trigger pulse on its strobe output pin at the start of each frame acquisition. The pulse triggers the analog output of a data acquisition card (NI-USB 6211, National Instruments, USA) which has a 16-bit, 250 kS/s sampling rate to send a ramp voltage signal to the galvo-scanning mirror with some user-specified delay. For the experiments, an exposure of 30 μs, which corresponds to a slit-width of about 2.6 μm, was chosen.

### LS-CLE data acquisition and processing protocol

In terms of the preparation of the urology specimens, small cut-outs from freshly excised urology specimens were acquired from the histopathology laboratory in Charing Cross Hospital (London, UK) and snap-frozen at –80 °C. These specimens were obtained from consented patients using Imperial College London tissue bank’s ethical protocol following the R-19016 project, with ICHTB HTA license number 12275 and REC approval reference 17/WA/0161. The neoplastic tissue was obtained from macroscopically visible tumor sites. Prior to image acquisition by the proposed LS-CLE-fiber actuated system, the frozen cut-outs were thawed at room temperature for 5 minutes. Following this, the samples were first immersed in test tubes containing acriflavine hydrochloride solution at 0.1% concentration. The fluorescent agent was left to stain the tissues for 30 seconds and then gently rinsed with water to wash off excess agent before imaging. The flexible fiber-bundle probe was autonomously steered at its distal end along a preoptimized spiral path onto the tissue surface, and images were obtained in real-time. At the end of the imaging process, the excess fluorescent agent was again wiped off the tissue surface.

A LabVIEW-based graphical user interface (GUI) was developed to acquire and record LS-CLE images at 120 fps. Endomicroscopy video loops were recorded for about 60 seconds from 4 to 6 sites on the tissue surface. The data were post-processed to remove the fiber-bundle *pixelation* artifacts (caused by the fiber cores) by convolution with a 2D Gaussian filter (sigma = 1.6 pixels, 1.4 μm on the bundle). This was followed by mosaicking the video frames (i.e., stitching overlapping frames) to produce images with a larger field of view (FOV) while still maintaining microscopic resolution that could be comparable to histology slides. A fast normalized cross-correlation (NCC) algorithm was used for pair-wise image registration (*70*). The primary intent behind using the fast NCC was to use Fast Fourier transform (FFT) to evaluate, in one pass, the correlation coefficient between consecutive image frames *I*_*k*_ and *I*_*k* +1_. A 2D correlation map was generated using:

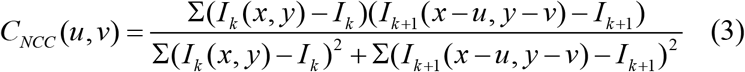

where *I*_*k*_ is the average pixel value of the image *K*; *x, y* are pixel coordinates, and *u, v* is the translational shift. The translational shift was obtained from the location of the maximum correlation peak in the 2D correlation map, given by:

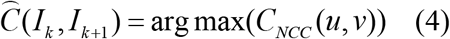

As the final step, a weighted-average feathering function for image blending was implemented to reconstruct an output mosaic with minimum visible seams.

### In vivo animal testing setup

A female Yorkshire swine of 45 kg was used for our animal testing. All procedures were conducted in accordance with the protocol approved by the Acibadem Ali Aydinlar University Animal Experiment Local Ethics Committee. The trial commenced by premedicating the swine with a combination of ketamine (preanesthetic and analgesic drug) and xylazine (sedative and analgesic drug with some muscle relaxation properties). After inducing anesthesia using propofol (anesthetic drug) through an intravenous cannula (inserted into the swine’s left ear vein), the swine was endotracheally intubated. During the trial, isoflurane (anesthetic drug) was used to maintain anesthesia, and metamizole (pain killer) to treat discomfort. Using the anesthesia machine (Leon Plus, Löwenstein Medical SE & Co. KG, Germany), the swine’s ventilation was maintained using mechanical ventilation under the continuous monitoring of its physiological status. At the end of this study, to facilitate euthanasia, the isoflurane dose was increased to a maximum (5%), and potassium chloride was administered intravenously after a 5-minute interval.

### Histopathological Assessment

The specimens from the liver and spleen were placed in an aqueous solution of formaldehyde and sent to an independent laboratory (Acibadem pathology/Nisantasi Pathology Group, Istanbul) for histopathological assessment. First, the tissue specimens were macroscopically examined. Subsequently, the tissue samples were serially sectioned, and all tissue planes were mapped and sampled. After embedding the tissue samples with paraffin, the samples were cut into 4μm thick slices and stained with hematoxylin and eosin (H&E). The H&E slides (Fig. 8E) were then assessed by an expert pathologist to investigate if there were any signs of thermal injury.

## Acknowledgments

MEMKA and JZ thank Dr. Giulio Dagnino for supporting the development of the computational interface for the microscopic image tracking system. MEMKA and JZ also thank Dr. Weibang Bai for his supportive role during the Da Vinci® surgical robot integration. HL would like to acknowledge the support of the Inha university research grant. MEMKA and BT thank Dr. Salzitsa Anastasova for her fruitful discussion on using *ex vivo* tissue samples and Dr. Meysam Keshavarz for his insight on the micromanipulation demonstrations. MEMKA would also like to acknowledge Andreas Leber and Prof. Fabien Sorin for their support with the rheology measurements. MEMKA and BT also thank Ms. Pelin Erkasap, Dr. Murat Saruç, and the Acibdaem CASE team for their support during the animal trials. MEMKA and BT acknowledge Herdeks Ltd team (Mr. Yusuf Akca, Mr. Alperen Kartal) for their support with the surgical laser system. Finally, MEMKA thanks Jiwoo Choi for providing support with videography.

## Author contributions

Conceptualization: MEMKA, HL, BT

Methodology: MEMKA, JZ, BR, DS, HK, KV, HL, YL, AAD, HU, GG, AA, ME, GY

Investigation: MEMKA, JZ, BR, BL, YL

Visualization: MEMKA, JZ, HK, RD, ZT

Funding acquisition: GY, EY, BT

Project administration: GY, EY, BT

Supervision: GY, EY BT

Writing – original draft: MEMKA, JZ, BR, BL

Writing – review & editing: GY, EY, RD, BT

## Data and materials availability

All data are available in the main text or the supplementary materials.

## Supplementary Materials

Materials and Methods

Supplementary Text

Figs. S1 to S14

References *(69 - 71)*

Movies S1 to S10

