## Supplemental File for "Fiberbots: Robotic fibers for high-precision minimally invasive surgery"

**This PDF file includes:**

Supplementary Text: Supplementary notes 1-5
Figs. S1 to S14
Captions for Movies S1 to S10
References *(71 - 73)*

**Other Supplementary Materials for this manuscript include the following:**

Movies S1 to S10

### Supplementary Text

#### Supplementary Note 1: Electronic circuitry for fiber actuation and sensing

The fiber was actuated using an electronic circuit made with four independent voltage channels, which impose an electrical current "i" to carry out the motion in four directions (X-, X+, Y- and Y+). This configuration was realized using two voltage-to-current converters built around power amplifiers (OPA548, Texas Instruments, USA) mounted in a non-inverting topology. By imposing a voltage difference at the output of each amplifier ($V_0^{out}$ and $V_1^{out}$, respectively), a current flow that is inversely proportional to the resistance of the wire conduction path (< 100 Ω), was generated through the single pair of wires running on the same side of the fiber. Twisting and connecting the distal ends of the pair of wires guaranteed a closed-loop for current circulation inside the fiber. By considering the voltage signals ($V_0^{in}$ and $V_1^{in}$) produced by the two channels inside a digital-to-analog (DAC) conversion board (NI-9264 DAC, National Instruments, USA) docked into a real-time controller (CompactRIO cRIO-9025, National Instruments, USA), and the equivalent resistance value for the respective wire-pair ($R_e$), the current was calculated as follows:

$$i = \frac{(V_0^{out} - V_1^{out})}{R_e} = \frac{2(V_0^{in} - V_1^{in})}{R_W + R_W + R_S} \approx \frac{(V_0^{in} - V_1^{in})}{R_W} \quad \text{(SI-1)}$$

$R_S$ was considered close to 0 Ω due to the high conductivity of the silver adhesive glue paint (CW2205, Chemtronics, USA) employed to bond each pair of wires, whereas $R_W$ was assumed similar between the wire pair. Irrespective of the direction of current flow, the electric power dissipated by one wire pair is equivalent to the quadratic factor presented in equation SI-2.

$$P_{electric} = i^2 R_e = 2R_W \frac{(V_0^{in} - V_1^{in})^2}{R_W^2} = 2\frac{(V_0^{in} - V_1^{in})^2}{R_W} \quad \text{(SI-2)}$$

An additional relay (G6K-2P-Y, Omron, Japan) activated by an emitter-follower transistor topology (2N2222, Multicomp Pro, USA) was intercalated between each current loop to form an automatic switch that disconnects the fiber from the electronics in the event of an emergency.

The circuit was converted into a printed circuit board (PCB) using a PCB design and electrical schematic software (EAGLE, Autodesk, USA) as part of a larger electronic assembly that includes the PCBs for power and temperature feedback measurements, as shown in **figure S4**. The bottom PCB in the assembly holds the electronics responsible for driving eight voltage-to-current converters (channels), whereas the middle PCB contains the proximal end of the electrothermal fiber with the internal wires soldered to the exposed electrical pads. Additional ultraviolet (UV)-curable adhesive (3525, Loctite, Germany) was poured above these soldering anchors to protect the wires and ensure mechanical stability to the fiber during actuation.

Power measurements were performed within the middle PCB by two electronic channels that measure, separately, the electric current flowing through one pair of wires and the voltage registered between the proximal ends (or input terminals) of each wire (**Fig. S4B**). Although a theoretical formula for the generated power was derived previously, in practice, some signal drifts, interferences, and changes in wire resistance during actuation lead to fluctuations in power level.

So, more accurate measurements were needed to better characterize the fibers. In order to achieve this, three amplifiers (OPA604, Texas Instruments, USA) in a buffer circuit topology were intercalated in sequence along the current loop formed by each pair of wires to produce two voltage measurements $(V_{P1} - V_{P2})$ and $(V_{P2} - V_{P3})$. Two amplifiers (Amp1 and Amp2) sense the voltage drop across a fixed-value resistor (10 Ω) in the loop, which enabled us to calculate the magnitude of the circulating current as shown below:

$$i = \frac{2(V_{P1} - V_{P2})}{10\ \Omega} \quad \text{(SI-3)}$$

Similarly, the voltage originated at the terminals of the pair of wires is detected by the two amplifiers (Amp2 and Amp3) and calculated according to the equation below:

$$V = 2(V_{P2} - V_{P3}) \quad \text{(SI-4)}$$

The voltage measurements $(V_{P1}, V_{P2}, V_{P3})$ were acquired using an analog-to-digital (ADC) converter (NI-9205 ADC, National Instruments, USA) docked into another slot of the same real-time controller (cRIO-9025). The measured power was computed as follows:

$$P_{meas} = i \cdot V = \frac{4(V_{P1} - V_{P2})(V_{P2} - V_{P3})}{10\ \Omega} \quad \text{(SI-5)}$$

Furthermore, the resistance of the wire pair was also estimated using the equation below, thus, allowing real-time feedback measurements for both power and resistance during fiber actuation.

$$R_e = \frac{P_{meas}}{i^2} = \frac{(V_{P2} - V_{P3})}{(V_{P1} - V_{P2})} \cdot 10\Omega \quad \text{(SI-6)}$$

In terms of temperature feedback measurements, four K-type thermocouples (406-590, TC Direct, UK) were placed along the outer circumference of the fiber next to the pair of wires (**Fig. 3R**), with 90° separation between them. Polyimide film tape (5413, 3M, USA) was used to hold the thermocouples at the measurement junction. Each thermocouple was then connected to an instrumentation amplifier (AD8497, Analog Devices, USA) at the reference junction to detect the voltage levels induced by the temperature variations (5 mV/°C), as shown in **fig. S4D**. The voltage measurements $(V_T)$ were acquired using the same ADC converter (NI-9205 ADC). The temperature was computed using the equation below:

$$T = \frac{V_T}{(5\ mV/^{o}C)} \quad \text{(SI-7)}$$

Supplementary Note 2: Outer surface characterization

As the surface temperatures of the actuated fiber are above the biological safe limit of 43°C (**Fig. 1H and Fig. 3T**), we introduced cooling by injecting compressed air through the central channel. We placed the fiber alongside the laser displacement sensor (LK-G5000, Keyence, Japan) inside an incubating mini shaker (VWR® Incubating Mini Shaker, VWR, UK) (**Fig. 3A**). The incubating mini shaker was used to set the environment temperature of the fiber to 38°C ± 1°C. A 21-gauge needle (SA7524, Adhesive Dispensing Ltd, UK) was connected from its hub to a 465 cm long air hose assembly (350 cm long polyethylene tubing (126-3103, RS, UK) connected to a 115 cm long flexible PVC tubing (WZ-30526-18, Cole-Parmer, USA)). The shaft of the needle, on the other hand, was inserted into the central channel of the fiber. Two K-type thermocouples (406-590, TC Direct, UK) were placed along the outer circumference of the fiber next to the pair of wires to measure the outer surface temperature. Our experiments show that using air at room temperature; we can decrease the surface temperature of the actuated side by approximately 45.5°C, down to 43°C. However, the temperature gradient between the opposing ends dropped from 24.2°C to approximately 9.2°C at 5 bar, and fiber tip displacement decreased by 1.94 mm. Using a fiber with an outer diameter of 1.65 mm and 11 cm in length, the maximum fiber tip displacement to avoid thermal tissue damage is 1.68 mm for an input power of 1 W (**Fig. 3B**).

Supplementary Note 3: Simplified analytical model of the electrothermally actuated fiber (referred to as fiberbot in the remainder of the text)

A simplified analytical model was developed to estimate the fiber tip displacement as a function of input power. Inspired by the model developed by Mirvakili et al. (*6*), the complex cross-sectional geometry of the electrothermal fiber was simplified into a bimorph structure as shown in **fig. S14A**, where the outer surface temperature of the actuated hot side of the structure is represented by $T_1$, and the outer surface temperature of the passive cold side of the structure is represented by $T_2$. Using this model, the cantilever bending of the structure is schematically represented in **fig. S14B**, where A is the amplitude of the fiber tip displacement, $L_0$ is the length of the neutral axis of the fiber, $L_{top}$ is the arc length of the actuated side of the fiber, $L_{bottom}$ is the arc length of the passive side of the fiber, $r_0$ is the radius of curvature with respect to the fiber's neutral axis $L_0$, $r_{top}$ is the radius of curvature with respect to the extended side of the fiber $L_{top}$ , $r_{bottom}$ is the radius of curvature with respect to the contracted side of the fiber $L_{bottom}$, W is the thickness of the fiber (i.e. outer diameter of the fiber).

Thermal strains are strains developed when a material is subjected to heating. In this model, we only consider the thermal expansion along the longitudinal direction of the structure. The one-dimensional thermal strain can be calculated as follows,

$$\varepsilon_T = \alpha_T \cdot \Delta T = \frac{\Delta L}{L_0} \quad \left(\Delta T = T_1 - T_2, \ \Delta L = L_{top} - L_{bottom}\right) \quad \text{(SI-12)}$$

where ε represents the thermal strain and $\alpha_T$ is the temperature-dependent coefficient of thermal expansion. The length and length change of the fiber can be obtained using the following equations:

$$L_0 = \theta \cdot r_0 \quad \text{(SI-13)}$$

$$\Delta L = L_{top} - L_{bottom} = \theta \cdot r_{top} - \theta \cdot r_{bottom} = \theta \cdot W \quad \text{(SI-14)}$$

where θ is the degree of curvature. By combining equations SI-12 and SI-14, and re-arranging, we get the following equation:

$$\varepsilon_T = \frac{\Delta L}{L_0} = \frac{\theta \cdot W}{\theta \cdot r_0} = \frac{W}{r_0} \Leftrightarrow r_0 = \frac{W}{\varepsilon_T} = \frac{W}{\alpha_T \cdot \Delta T} \quad \text{(SI-15)}$$

Using the Pythagorean theorem, the radius $r_0$ can be calculated as follows:

$$r_0 = \sqrt{(r_0 - A)^2 + L^2} \Leftrightarrow r_0 = \frac{A^2 + L^2}{2A} \approx \frac{L^2}{2A} \quad (A \ll L) \quad \text{(SI-16)}$$

The amplitude of the fiber tip displacement (A) can be computed as shown below:

$$A \approx \frac{\alpha_T \cdot \Delta T \cdot L^2}{2W} \approx \alpha_T \cdot \frac{\Delta T \cdot L_0^2}{2W} \quad \text{(SI-17)}$$

Thus, the displacement of the fiber is proportional to the average temperature difference between the opposing sides of the fiber. In turn, this temperature difference arises from the heat generated by Joule's effect that matches the electric power carried by each stainless-steel wire inside the fiber, transferred to the PC body of the fiber by thermal conductivity with minimal loss to the surrounding environment (by convection).

If we consider the actuated hot side of the fibre at a steady state, the following formula can be derived according to the law of energy conservation:

$$\dot{Q}_{Joule} - \left(\dot{Q}_{rad\_A} + \dot{Q}_{conv\_A} + \dot{Q}_{cond}\right) = 0 \quad \text{(SI-18)}$$

where $\dot{Q}_{Joule}$ represents the heat generated by the Joule's effect. $\dot{Q}_{rad\_A}$ represents the rate of heat transfer by emitted radiation to the environment (represented by temperature $T_e$) from the actuated side. $\dot{Q}_{conv\_A}$ represents the rate of heat transfer by convection to the environment from the actuated side. $\dot{Q}_{cond}$ represents the heat transfer rate from the actuated (high) temperature side to the passive (low) temperature side of the fiber by conduction.

$$\dot{Q}_{Joule} - \left(\varepsilon \cdot \frac{A_e}{2} \cdot \sigma \cdot ({T_1}^4 - {T_e}^4) + h_{conv} \cdot \frac{A_e}{2} \cdot (T_1 - T_e) + K \cdot A_{cond} \cdot \frac{dT}{dx}\right) = 0 \quad \text{(SI-19)}$$

Equation SI-19 can be rewritten as follows:

$$\varepsilon \cdot \frac{A_e}{2} \cdot \sigma \cdot ({T_1}^4 - {T_e}^4) + h_{conv} \cdot \frac{A_e}{2} \cdot (T_1 - T_e) = \dot{Q}_{Joule} - K \cdot A_{cond} \cdot \frac{dT}{dx} \quad \text{(SI-20)}$$

where ε is the emissivity of the encapsulating polymer, σ is the Stefan-Boltzman constant ($5.67 \times 10^{-8}\, {W}/{m^2} \cdot K^4$). $T_e$ is the environment temperature, $h_{conv}$ is the convective heat transfer coefficient, $A_e$ is the surface area of the fiber in contact with the environment, K is the thermal conductivity of the polymer, and $A_{cond}$is the surface area of the overlapping surface between the hot actuated side of the fiber and the cold passive side of the fiber. $\dot{Q}_{rad\_A}$ can be rewritten as follows:

$$\varepsilon \cdot \frac{A_e}{2} \cdot \sigma \cdot ({T_1}^4 - {T_e}^4) = \varepsilon \cdot \frac{A_e}{2} \cdot \sigma \cdot ({T_1}^2 + {T_e}^2) \cdot (T_1 + T_e) \cdot (T_1 - T_e) \quad \text{(SI-21)}$$

Similarly, integrating the Fourier conduction equation ($\dot{Q}_{cond}$) results in:

$$K \cdot A_{cond} \cdot \frac{dT}{dx} = K \cdot A_{cond} \cdot \frac{\Delta T}{x1 - x2} \quad \text{(SI-22)}$$

In the case of the cold side of the fiber at a steady state, the following formula can be derived according to the law of energy conservation:

$$\dot{Q}_{cond} - (\dot{Q}_{rad\_P} + \dot{Q}_{conv\_P}) = 0 \quad \text{(SI-23)}$$

$\dot{Q}_{rad\_P}$ represents the rate of heat transfer by emitted radiation to the environment (represented by temperature $T_e$) from the passive side. $\dot{Q}_{conv\_P}$ represents the rate of heat transfer by convection to the environment from the passive side.

$$K \cdot A_{cond} \cdot \frac{\Delta T}{x1 - x2} - \left(\varepsilon \cdot \frac{A_e}{2} \cdot \sigma \cdot ({T_2}^4 - {T_e}^4) + h_{conv} \cdot \frac{A_e}{2} \cdot (T_2 - T_e)\right) = 0 \quad \text{(SI-24)}$$

Equation SI-24 can be rewritten as follows:

$$\varepsilon \cdot \frac{A_e}{2} \cdot \sigma \cdot ({T_2}^4 - {T_e}^4) + h_{conv} \cdot \frac{A_e}{2} \cdot (T_2 - T_e) = K \cdot A_{cond} \cdot \frac{\Delta T}{x1 - x2} \quad \text{(SI-25)}$$

Dividing equation SI-20 by equation SI-25 results in:

$$\frac{\varepsilon \cdot \frac{A_e}{2} \cdot \sigma \cdot ({T_1}^4 - {T_e}^4) + h_{conv} \cdot \frac{A_e}{2} \cdot (T_1 - T_e)}{\varepsilon \cdot \frac{A_e}{2} \cdot \sigma \cdot ({T_2}^4 - {T_e}^4) + h_{conv} \cdot \frac{A_e}{2} \cdot (T_2 - T_e)} = \frac{\dot{Q}_{Joule} - K \cdot A_{cond} \cdot \frac{\Delta T}{x1 - x2}}{K \cdot A_{cond} \cdot \frac{\Delta T}{x1 - x2}} \quad \text{(SI-26)}$$

Rewriting equation SI-26 gives:

$$\frac{\dot{Q}_{Joule}}{\Delta T} = \frac{\varepsilon \cdot \sigma \cdot ({T_1}^2 + {T_e}^2 \quad \cdot (T_1 + T_e) + h_{conv}}{\varepsilon \cdot \sigma \cdot ({T_2}^2 + {T_e}^2 \quad \cdot (T_2 + T_e) + h_{conv}} \cdot \frac{K \cdot A_{cond}}{x1 - x2} + \frac{K \cdot A_{cond}}{x1 - x2} \quad \text{(SI-27)}$$

Therefore, the electric power applied to the stainless-steel wires, which leads to $\dot{Q}_{Joule}$, can be approximated as a linear relationship with the temperature difference, and in turn, the displacement of the fiber tip (equation SI-17).

Supplementary Note 4: Numerical simulations of the fiberbots

To build upon the analytical model and further explore the temperature distribution and motion behavior of the fiberbot, we simulated the actuation principle by designing a three-dimensional (3D) model of the fiber embedded with stainless-steel wires using a finite element analysis (FEA) software (ANSYS R18.2, ANSYS Corp., USA). The finite element modules, including Thermal-Electric, Transient Thermal, and Static Structure analysis, were used to: (a) examine the effect of thermal expansion on the polymer-metal composite, and (b) assess different fiber cross-sectional structures.

We first investigated a solid polymer fiber body with two stainless-steel wires on one side as shown in **fig. S1 (A to E)**. The thickness of the fiber was 1.65 mm and the length of the fiber is 10 cm. The fiber body is defined as PC (density = 1.2 g/cm$^3$, coefficient of thermal expansion = 2.1 x $10^{-5}$ °C$^{-1}$, Young's modulus = 2 x $10^9$ Pa, Poisson's ratio = 0.42, isotropic thermal conductivity = 0.22 W/(m.°C), specific heat = 1200 J/(kg. °C)), whereas the wires were defined as stainless-steel (density = 7.5 g/cm$^3$, coefficient of thermal expansion = 1.04 x $10^{-5}$ °C$^{-1}$, Young's modulus = 1.9 x $10^{11}$ Pa, Poisson's ratio = 0.265, isotropic thermal conductivity = 25 W/(m.°C), specific heat = 490 J/(kg.°C), isotropic resistivity = 6.9 x $10^{-7}$ Ω.m). In the simulation environment, we employed the sweep mesh method to generate the 3D computational mesh for the fiber structure. The sweep mesh method was used to maintain high solver accuracy while reducing the mesh cell elements, leading to quicker solve times. The 3D fiber model was divided into 20 elements in the axial direction, and triangular elements with an edge resolution of 0.04 mm were used to mesh the cross-section. For the Thermal-Electric analysis module, we applied a potential difference of 6 V across two stainless-steel wires and obtained the Joule heat generated rate, which was equivalent to 0.512 W. The power generated by Joule heating was then imported into the Transient Thermal module and used to set the wires as a heat source at the beginning of the simulation time. The convection heat transfer coefficient (2.5 x $10^{-5}$ W/(mm$^2$·°C)) and emissivity (0.92) were set at the outer surface of the fiber.

From the Transient Thermal analysis, we obtained the temperature distribution of the fiber as exhibited in **fig. S1B**. The temperatures at the opposing sides (i.e., actuated side and passive side) of the fiber and the difference between them are shown in **fig. S1C**. The maximum temperature on the surface was 66.8°C, and the temperature difference across the fiber was 12.8°C. The temperature of the encapsulating polymer and the stainless-steel wires were imported into the Static Structure analysis module to calculate the deflection of the fiber. Under this module, the fiber model was fixed at one end, while the other end was allowed to deform freely in space (resembling a cantilever structure), with no-slip boundary conditions imposed at the interface between the metal and polymer. By actuating the pair of wires, one-directional deformation of the fiber was achieved, as shown in **fig. S1D**. The displacement of the fiber shown in **fig. S1E**, increased to a maximum of 0.587 mm during the first 4 seconds of the simulation and then decreased to 0.057 mm (70 seconds after actuation). The observed reversing of the motion is suspected to be due to the stainless-steel wires having a smaller coefficient of thermal expansion than the polymer. As the temperatures of both sides are elevated, the cold side expands faster than the hot side, whose expansion is restricted by the wires, causing a reduction in deflection. The calculated stress on the stainless-steel wires in **fig. S1E** demonstrates the force building up between the polymer and the wires with the increasing temperature, limiting and reversing the bending motion of the fiber.

A second pair of stainless-steel wires were introduced to the opposite side of the fiber, as shown in **fig. S2 (A to E)**., which facilitated the actuation in the reverse direction and eliminated the undesired reduction in deflection by enforcing mechanical symmetry. From the Transient Thermal analysis, adding another pair of wires had a negligible influence (less than 0.2%) on the temperature distribution on the fiber, as shown in **fig. S2, B and C**. In the Static Structure analysis module, the displacement of the fiber was 0.707 mm (**fig. S2, D and E**), and as expected, the fiber displacement did not decrease after reaching the maximum displacement position.

The fiberbot embedded with four pairs of wires and a central channel is illustrated in **fig. S3 (A to E)**. While the four pairs of wires allow bending in multiple directions, the central channel was introduced to deliver surgical tools and create a thermal barrier across the fiber. The Transient Thermal analysis results in **fig. S3B** show the resulting temperature distribution in the presence of a central channel. An air domain (specific heat = 1003 J/(kg°C)) was defined inside the central channel of the fiber to simulate the real-case scenario. The steady-state temperature difference between the actuated and passive sides was 21.6°C (**fig. S3C**), an increase by a factor of ~1.7 when compared to the temperature difference obtained in the absence of a central channel. The Static Structure analysis module resulted in a tip displacement value of 1.197 mm (**fig. S3E**), with a similar increase in the range of motion by a factor of 1.7 due to the introduced channel.

Supplementary Note 5: Electrical safety of the fiberbots

Most living tissues are capable of carrying electrical currents in their physiological state, which can induce harmful effects to them depending on the magnitude (> 100 μA), duration and regime of current propagation (i.e., direct current – DC, or alternating current, AC). Stimulation of body tissues with DC for extended periods of time has been shown to cause the most severe physiological impairments due to the interaction of direct conductive currents with tissues, leading to electrolysis and permanent migration of ionic species, which may result in tissue bruises and also burn from the generated DC heat *(71)*. To evaluate the electrical safety of our fiberbot, we conducted an experimental test to measure the level of current leakage from the tip of the fibre while actuated (i.e., current circulating through a pair of wires) and immersed inside a conductivity solution of 0.1413 S/m (HI7030L, Hanna Instruments, USA), as shown in fig. S10. The reason behind the selection of this conductivity level is related to the typical values recorded amongst biological tissues in the chest/abdomen cavity (namely, small intestine: 0.164 S/m, stomach: 0.164 S/m and lungs: 0.101 S/m *(72)*) by impedance spectroscopy methods within the α- and β-dispersion region bands of Schwan's dispersion theory *(71, 73)*. If electrical current escapes the fiberbot, it will travel through the solution due to the difference in electrical potential set between an immersed metal electrode and the electronic ground of the actuation device (acting as a current sink or lower impedance point), thereby creating a closed-loop for current flow and detection by series intercalation of an ammeter (2100035, Farnell, UK). As a result of our proper electrical isolation applied at the tip of the fibrebot (achieved by recessing the internal wires and the application of cyanoacrylate), we were able to detect zero amperes of current from the monitor of the ammeter (set at the μA range, resolution of 0.01 μA) for several tested input powers in the range between 0.04 W and 2 W. Thereby asserting the electrical safety of the proposed fiberbot during actuation, as also observed during the *in vivo* trial with the porcine model.

### Figures

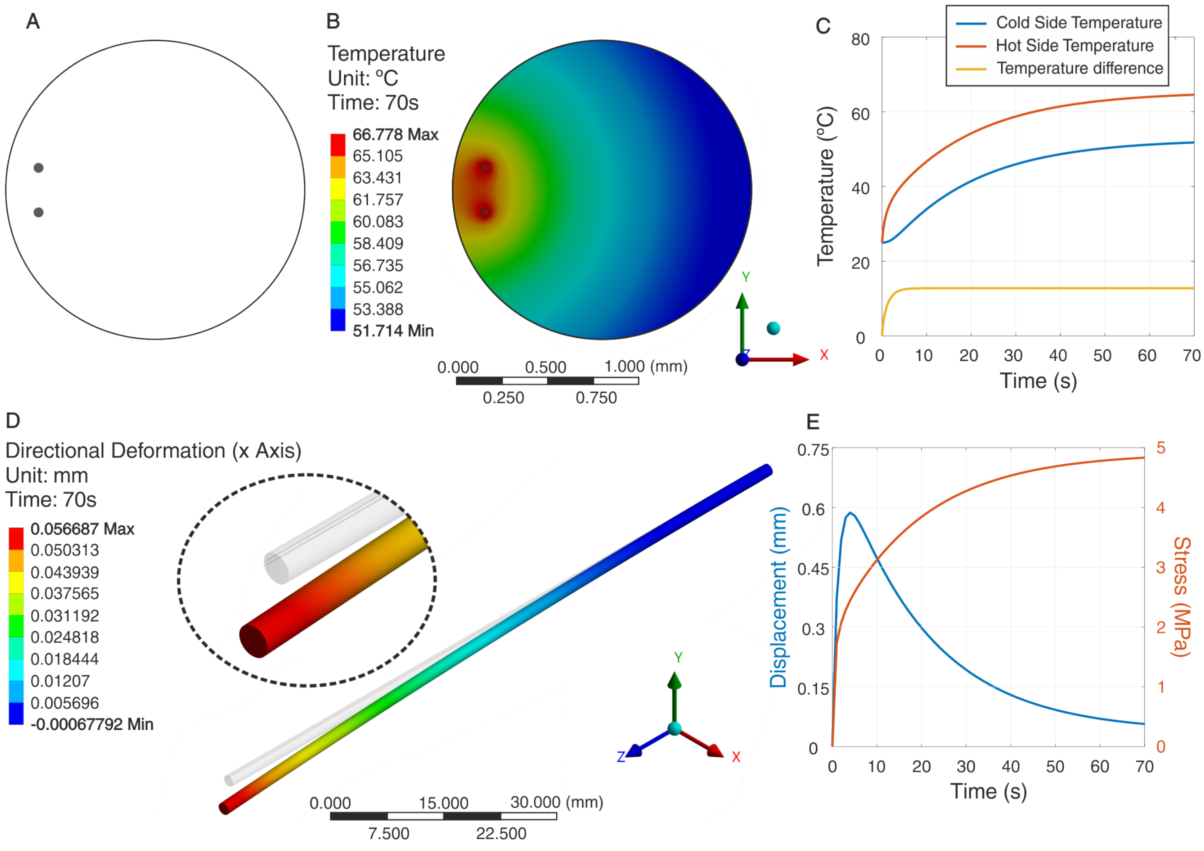


**Fig. S1. Finite element analysis of the two-wire fiberbot.** (**A**) 3D model of the one pair of wires cross-section. (**B**) Temperature distribution across the cross-section of the actuated fiber. (**C**) Temperature changes on the fiber's actuated and passive sides over time, and the temperature difference between them. (**D**) Directional deformation of the fiber as seen from the side view (length of the fiber). (**E**) Deformation change (tip displacement) induced by the simulated electrothermal effect and the highest mechanical stress level registered on the fiber.

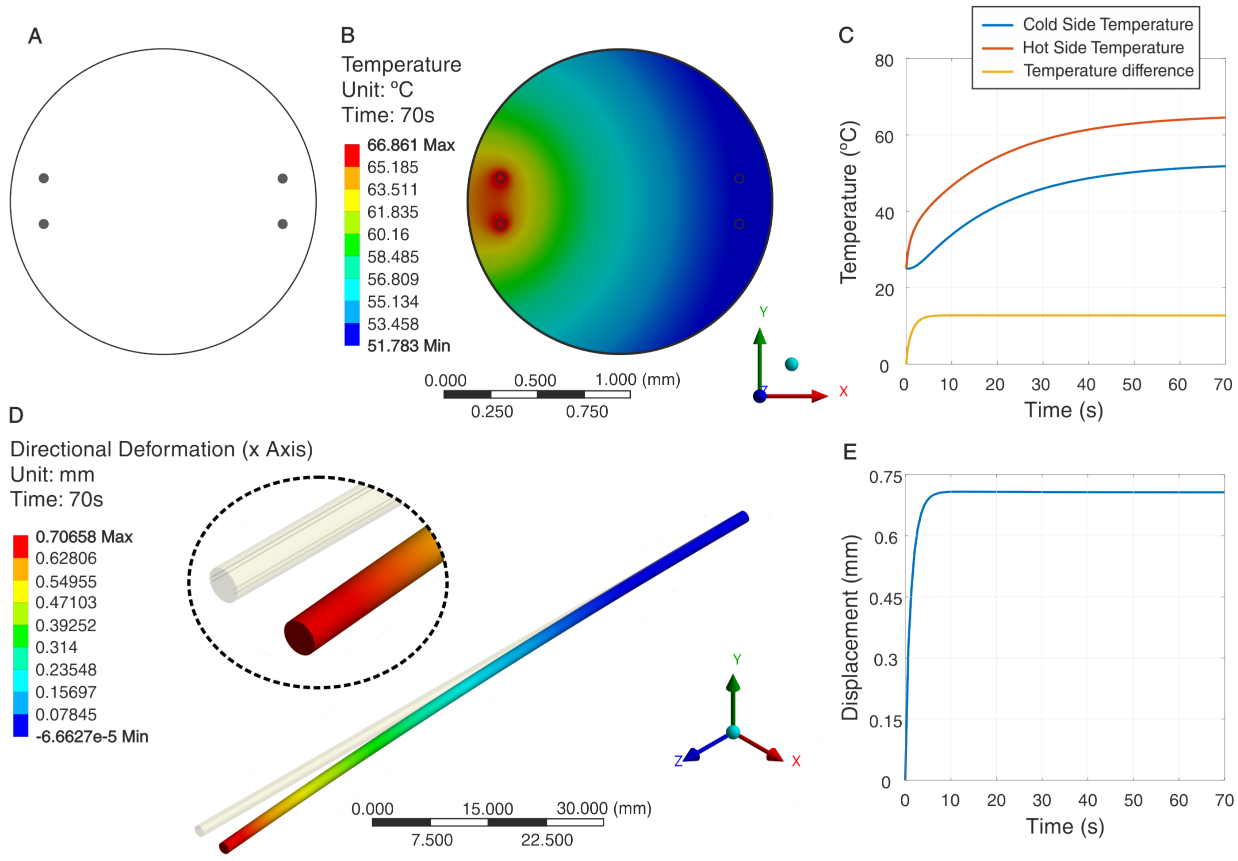


**Fig. S2. Finite element analysis of the four-wire fiberbot.** (**A**) 3D model of the two pairs of wires cross-section. (**B**) Temperature distribution across the cross-section of the actuated fiber. (**C**) Temperature changes on the fiber's actuated and passive sides over time, and the temperature difference between them. (**D**) Directional deformation of the fiber as seen from the side view (length of the fiber). (**E**) Deformation change (tip displacement) induced by the simulated electrothermal effect.

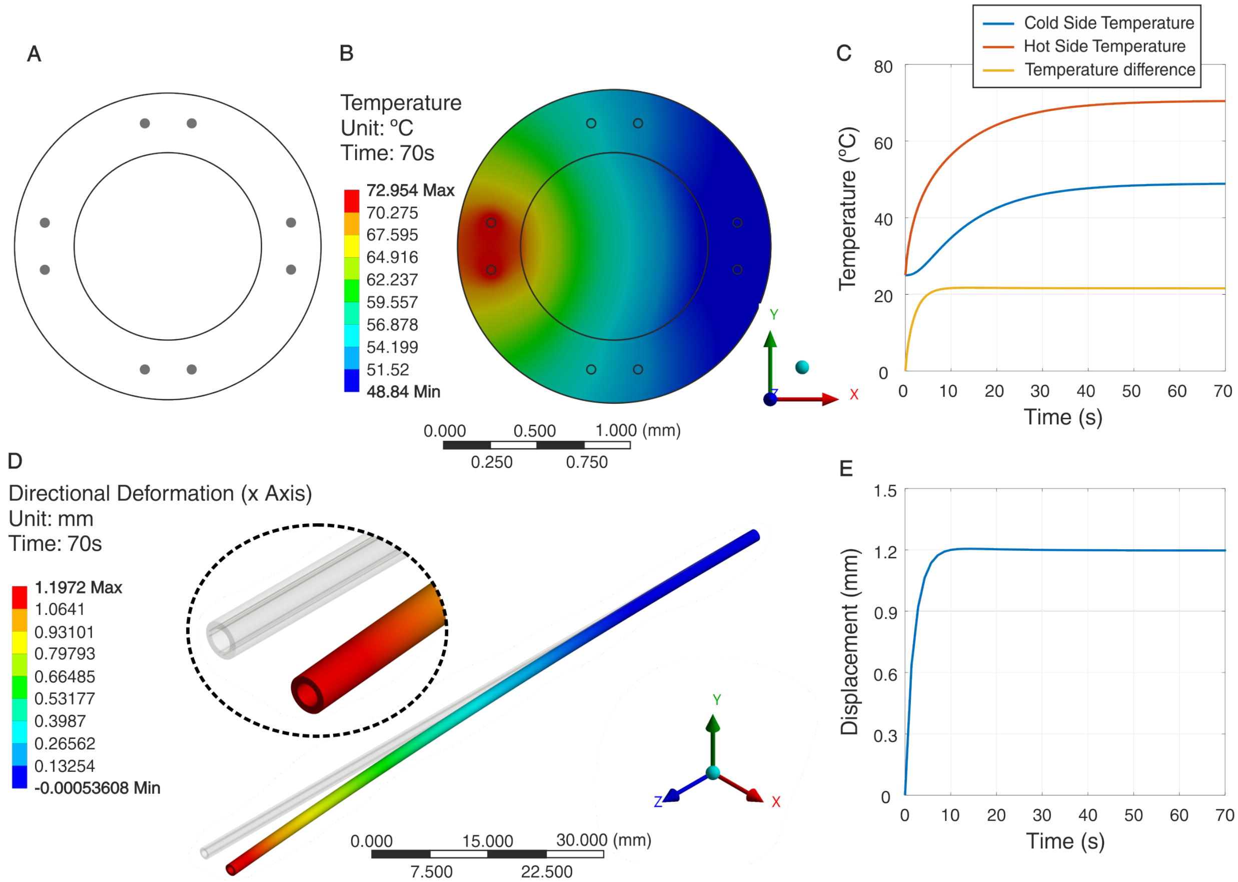


**Fig. S3. Finite element analysis of the eight-wire fiberbot.** (**A**) 3D model of the four pairs of wires cross-section. (**B**) Temperature distribution across the cross-section of the actuated fiber. (**C**) Temperature changes on the fiber's actuated and passive sides over time, and the temperature difference between them. (**D**) Directional deformation of the fiber as seen from the side view (length of the fiber). (**E**) Deformation change (tip displacement) induced by the simulated electrothermal effect.

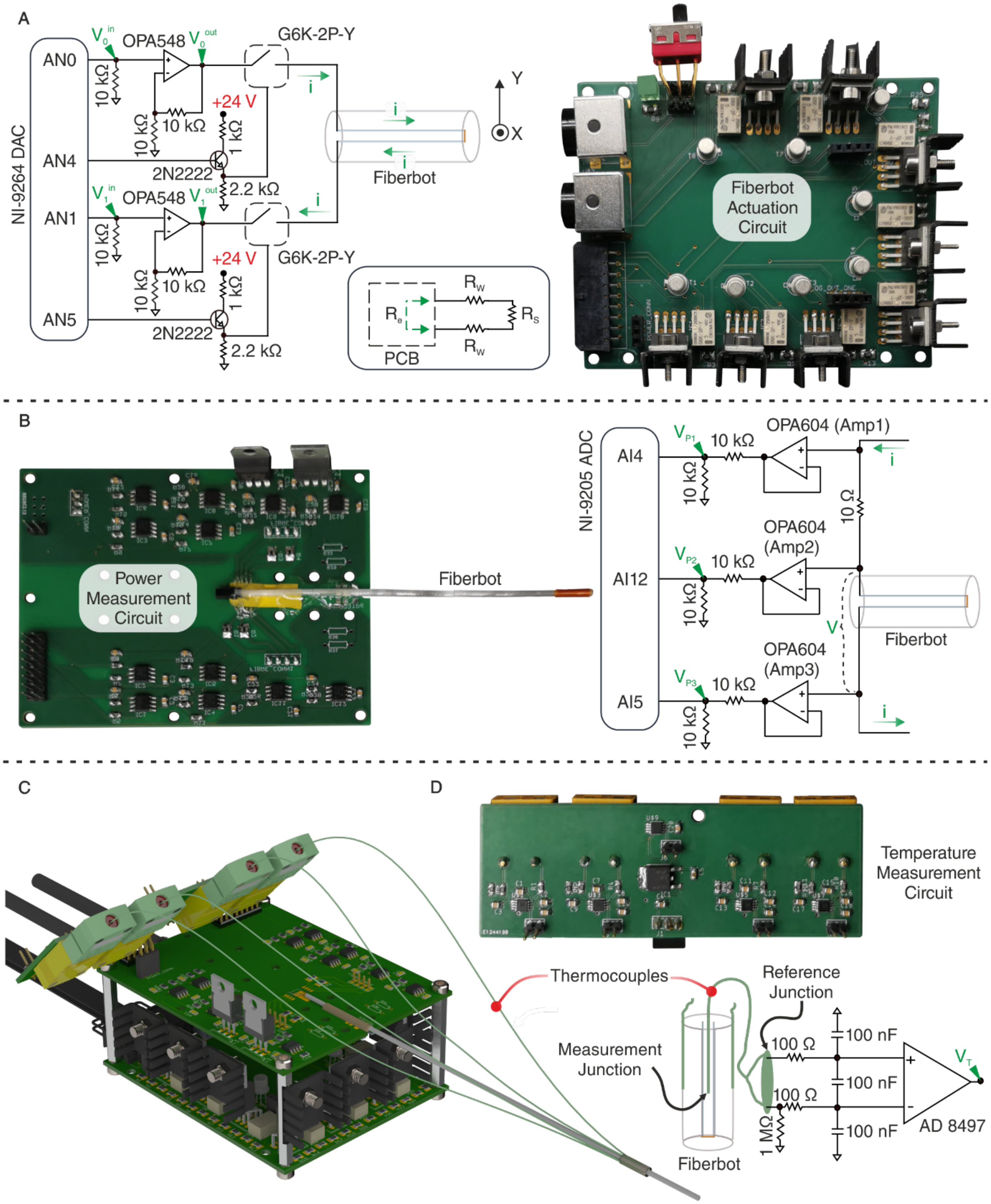


**Fig. S4. Electronic circuitry for fiberbot actuation and sensing.** (**A**) Simplified electronic schematic of the main components involved in generating the electrical current circulating through one wire-pair embedded inside the fiberbot (actuation). Inset: Equivalent resistance model of the wire pair. (**B**) Electronic schematic employed for real-time power feedback measurements. (**C**) Illustration of the assembled PCBs. (**D**) Electronic schematic employed to measure the temperatures utilizing external thermocouples placed around the outer surface of the fiber.

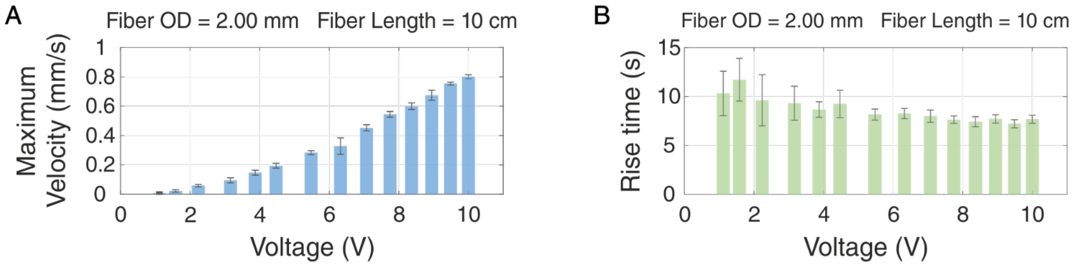


**Fig. S5. Maximum velocity and rise time of the fiberbot (derived from the characterization experiments that resulted in the velocity profiles presented in Figure 2L).** (**A**) Fiber's maximum velocity. (**B**) Average time required by the fiber to move from 10% to 90% of its fully developed steady-state positions, i.e., rise time for the different step input voltages.

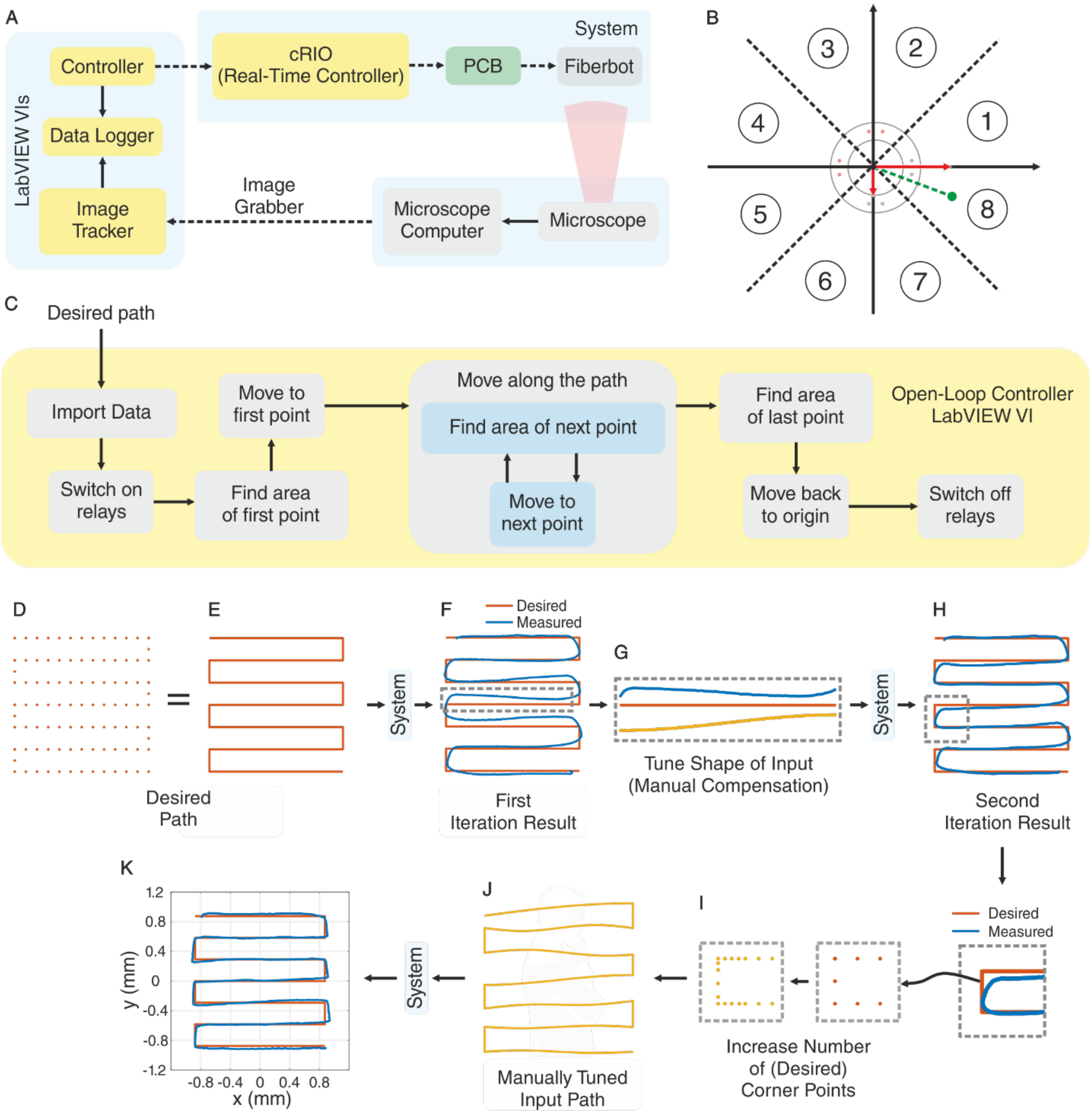


**Fig. S6. Open-loop control mechanism for fiberbot actuation**. (**A**) Schematic of the different modules, systems and graphical interfaces used to control the actuation process with microscope image tracking of the fiber tip. (**B**) Cross-sectional plane spanned by the fiber tip divided into 8 regions for accurate steerability along the different projection axes. (**C**) Flowchart of the implemented open-loop controller. (**D**), (**E**) Desired trajectory path (raster) formed by input coordinate points (2D) to the controller. (**F**) – (**H**) Positional differences between the desired and measured (image) paths observed after the first iteration of the open-loop controller, highlighting the shift in the horizontal line shape necessary for manual tuning of the input trajectory (second iteration). (**I**), (**J**) Additional tuning of the trajectory by manual increase in the number of coordinate points allocated to the trajectory corners. (**K**) Planar overlap of the desired and measured paths after manual optimization with the open-loop controller.

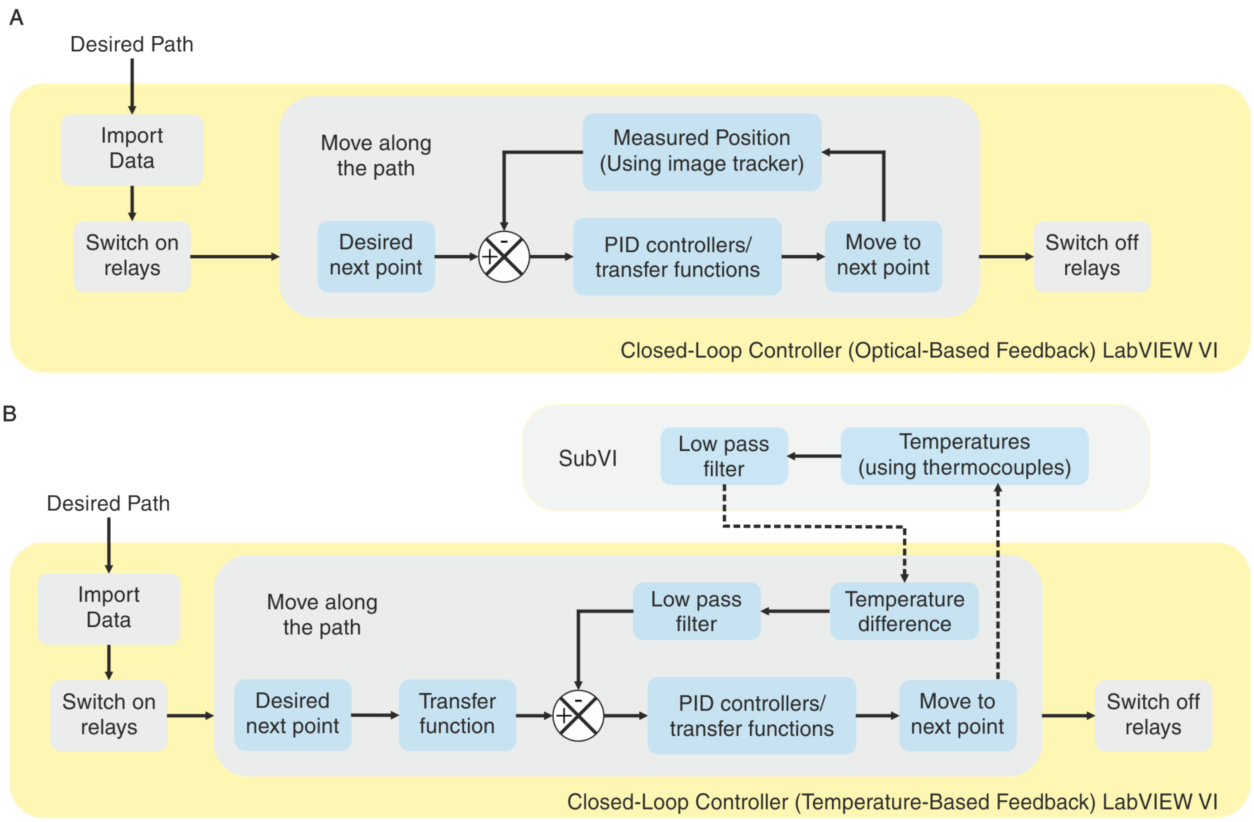


**Fig. S7. Closed-loop control mechanisms for fiberbot actuation.** (**A**) Flowchart of the control mechanism with microscope image feedback (measured tip position) used as input to the different PID controllers involved in the automatic estimation of the next trajectory point to be reached. (**B**) Flowchart of the control mechanism with temperature signal feedback obtained by the thermocouples placed on opposing sides of the actuated fiber and then fed to the PID controllers (low-pass filtering) to move to the next trajectory position.

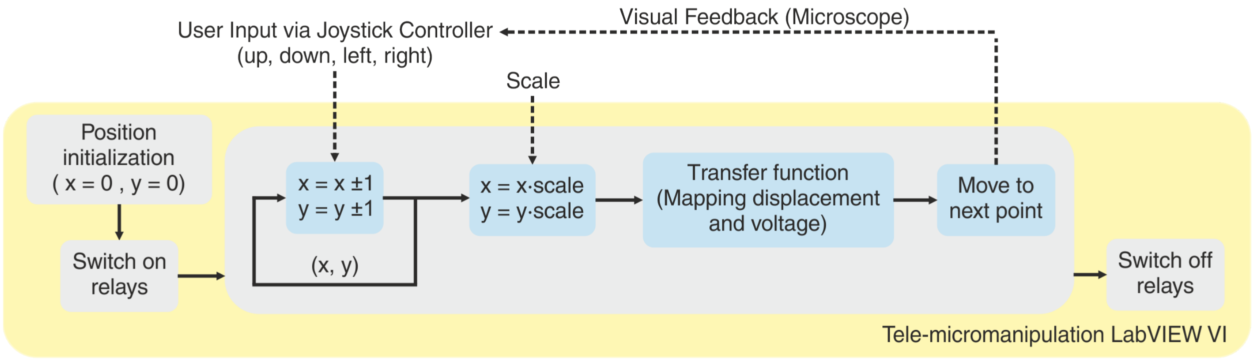


**Fig. S8. Flowchart of the implemented magnetic tele-micromanipulation system.**

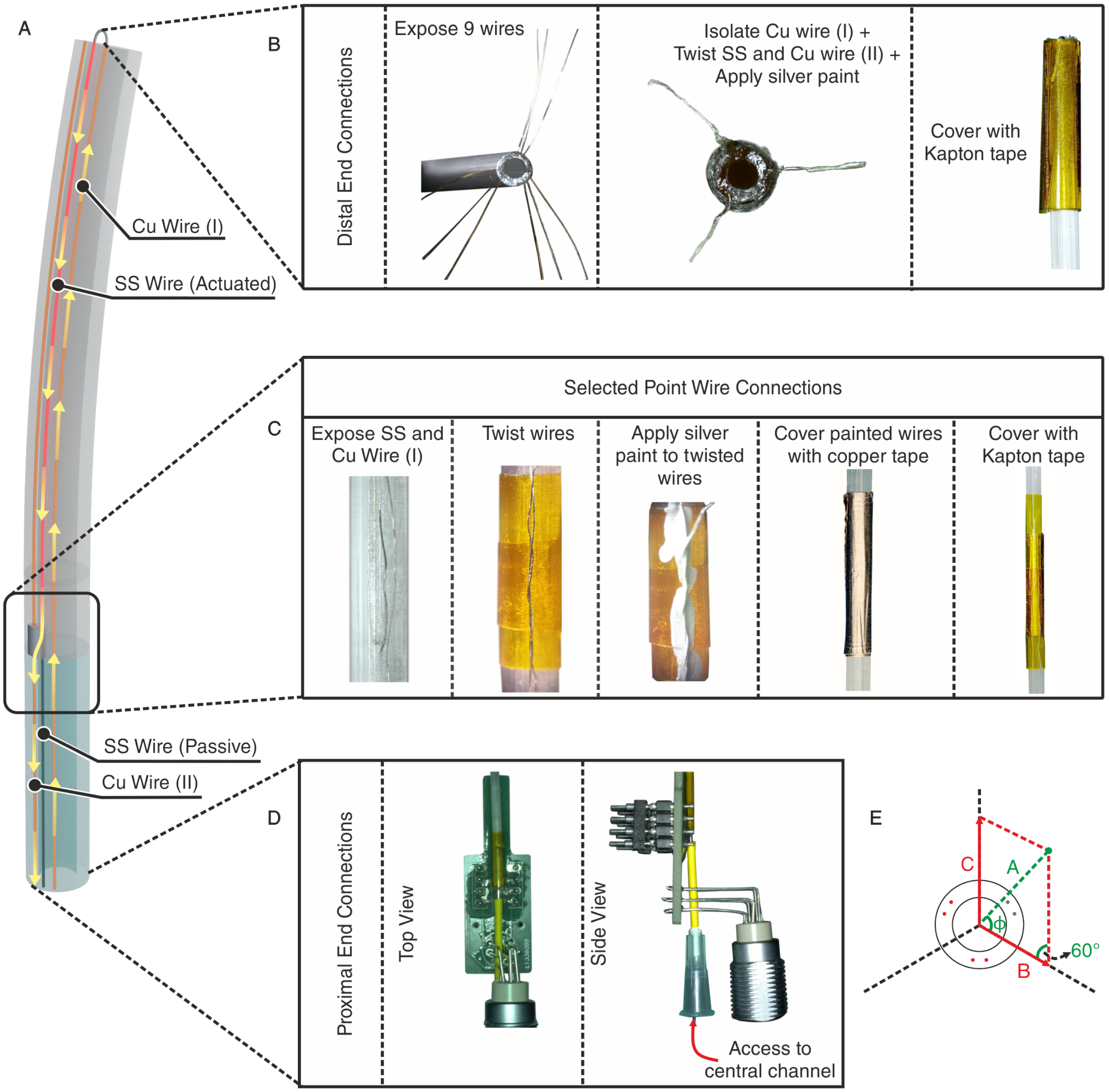


**Fig. S9. Selective fibrebot actuation.** (**A**) Schematic for the electric current circulation (arrows) through the active section of the fiber (copper – Cu – and stainless-steel – SS – wires) and coupling to the passive section (Cu wires only). (**B**) Detail of the distal tip of the active fiber section with 9 exposed wires (triplets of two Cu wires and a single SS wire) and electrical connection made to close the loop for electric current circulation, followed by protection from the exterior (Kapton tape). (**C**) Detail of the selected point connection over the fiber body that makes the separation between the (electrothermally) active and passive sections. Different steps involved in the exposure and final closure of the embedded wires in the electrical connection. (**D**) Detail of the proximal end connection of the fibre, which involves Cu wire soldering and fibre polymer anchoring to a circuit board with electric connections to the PCB actuation system and a central lumen access through a needle. (**E**) Coordinate system (and angles) employed for the estimation of the position of the fibre tip actuated by 3 pairs of wires.

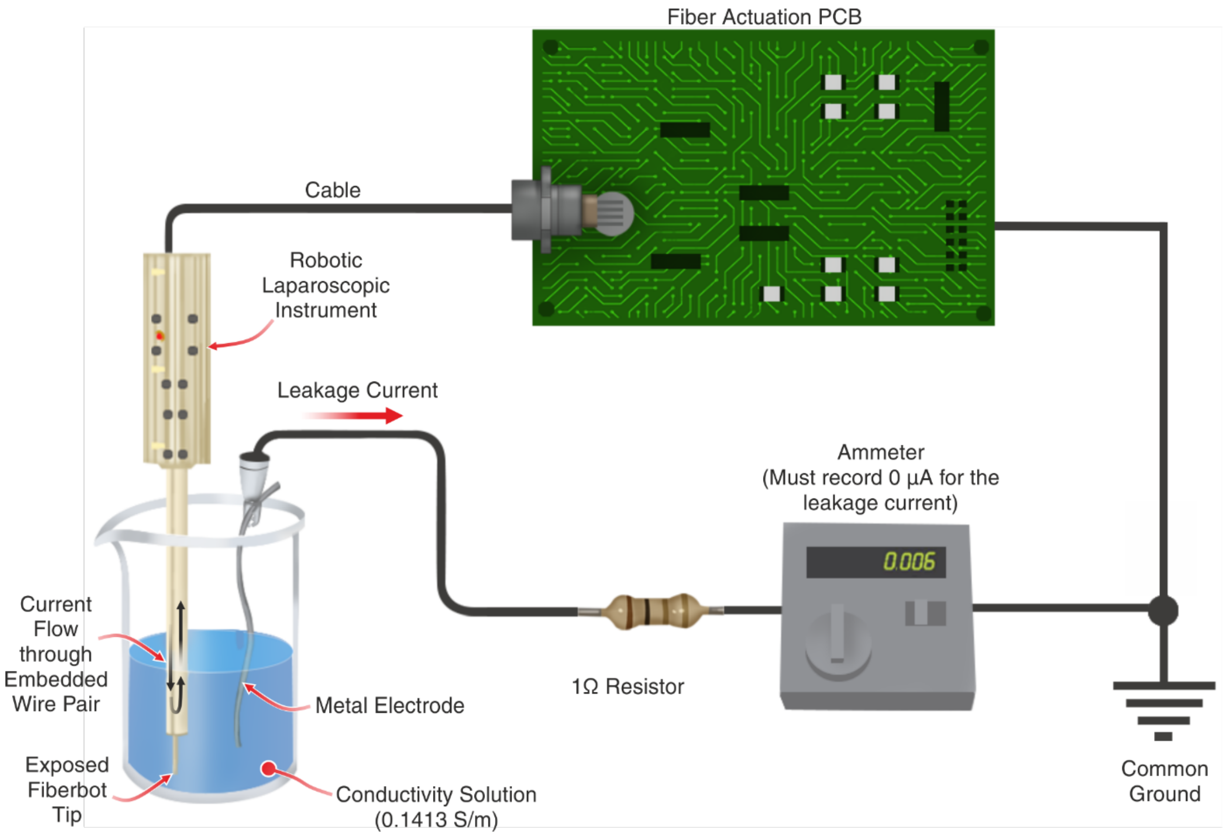


**Fig. S10. Setup developed to evaluate the electrical safety of the robotic laparoscopic instrument.**

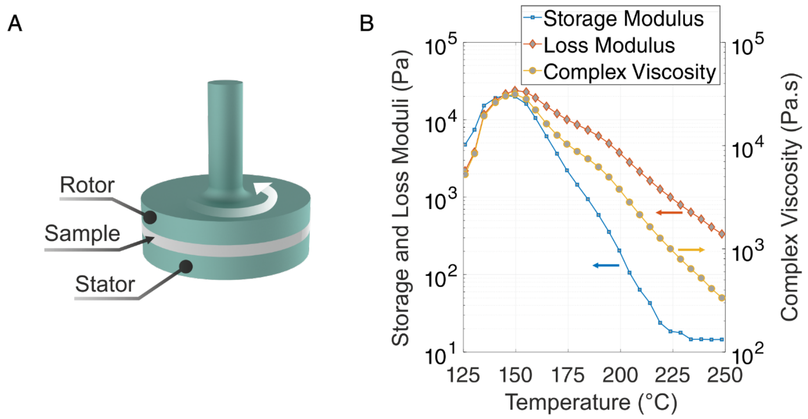


**Fig. S11. Rheological properties of 3D printed PC during a temperature ramp in oscillatory shear rheology.** (**A**) Schematic of the testing setup for measuring the storage modulus, loss modulus and complex viscosity of the 3D printed sample. (**B**) The complex viscosity decreases as a function of temperature. The loss modulus decreases and crosses over the decreasing storage modulus. The experiment was conducted using a rheometer with an environmental test chamber (AR2000ex, TA Instruments, USA) at an angular frequency of 1 rad/s and frequency of 0.159 Hz.

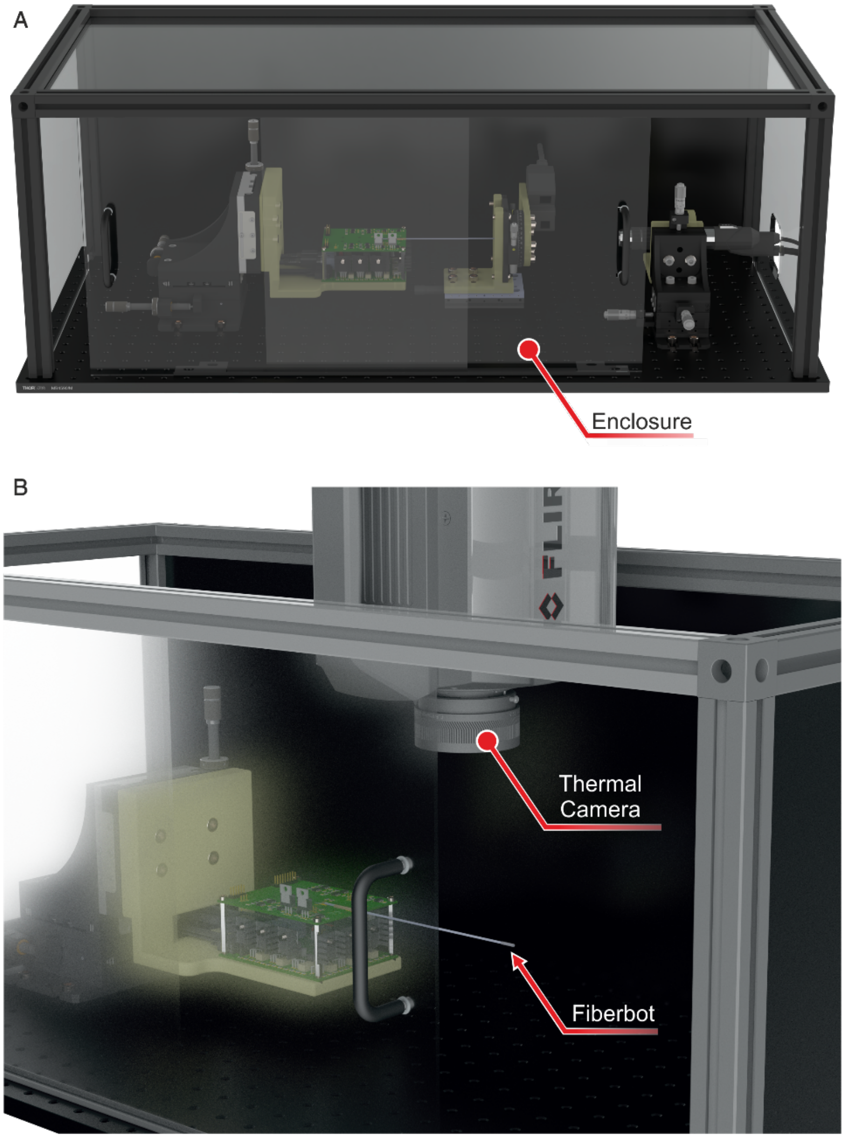


**Fig. S12. Setup developed for complete fiberbot motion characterization and coupling to other measurement systems.** (**A**) Enclosed case assembled to protect the electrothermal fiber from external environmental sources of light and airflow. (**B**) Positioning of the thermal camera above the fiber setup for free-range movement estimation with thermographic images.

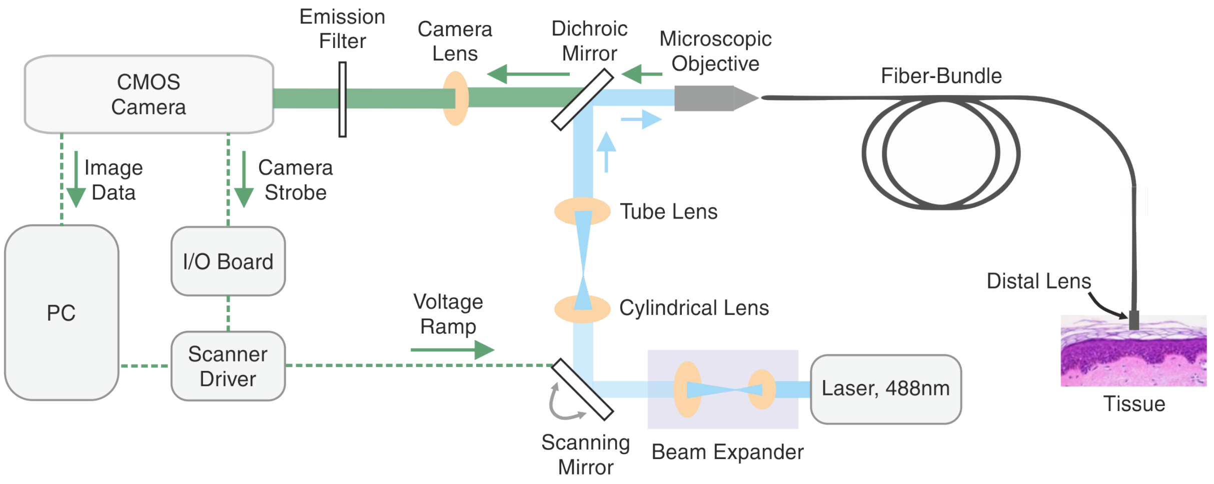


**Fig. S13. Line-scan confocal laser endomicroscopy (LS-CLE) system's schematic with main composing modules for light generation, splitting and detection coupled to a fibre-bundle probe for urothelial cancer imaging.**

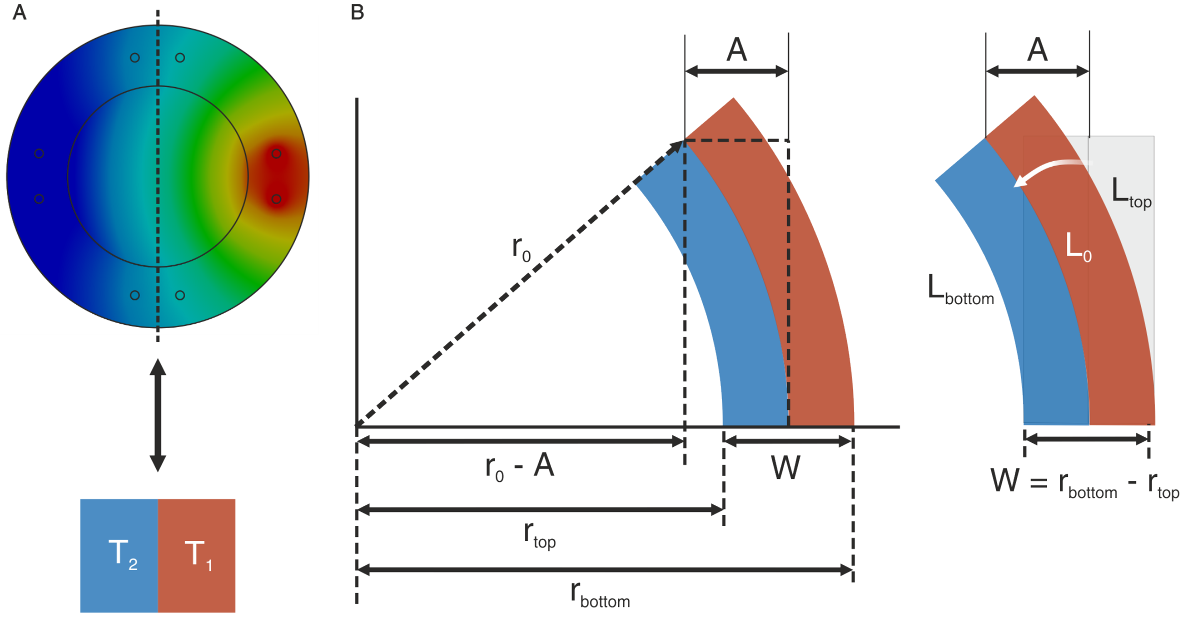


**Fig. S14. Simplified analytical model for electrothermal fiberbot actuation.** (**A**) Modelling of the generated heat along the cross-section of the fiber, approximated by two different temperature values symmetrically distributed on opposing sides of the fiber. (**B**) Schematic employed as reference for the one-dimensional bending model of the electrothermal fiber.

**Movies:**

**Movie S1.** First part: The unexpected movement of the thermally drawn fiberbot when it is placed on top of a hot plate due to asymmetric thermal expansion. Second part: The fiberbot moves along a circular path with three different actuation speeds and circles radii. Third part: The fiberbot moves along a circular path inside a scanning electron microscope. The fiberbot is 1.65 mm in outer diameter and 12 cm in length.

**Movie S2.** Numerical simulation for the temperature and mechanical displacement of the fiberbot when actuated through a single pair of wires (left side).

**Movie S3.** Longitudinal temperature distribution of the actuated fiberbot using a thermal imaging camera. Fiber is 1.65 mm in outer diameter and 12 cm in length.

**Movie S4.** Magnetic tele-micromanipulation of a magnetic object through two different tasks: (a) pick and place the object, moving from, e.g., A to B, and (b) move through a maze/labyrinth. The fiberbot is 1.65 mm in outer diameter and 12 cm in length.

**Movie S5.** Magnetic tele-micromanipulation of a magnetic object through two different tasks: (a) weave the object (ball) through football (soccer) cones, and (b) pass through defenders to score a goal. The fiberbot is 1.65 mm in outer diameter and 12 cm in length.

**Movie S6.** The fiberbot moves along a preoptimized spiral path while simultaneously imaging lens tissue and *ex vivo* bladder tissue using a fiber-bundle-based line-scan confocal laser endomicroscopy system. The fiberbot is 1.65 mm in outer diameter and 12 cm in length.

**Movie S7.** The fiberbot moves along a preoptimized path of parallel lines while simultaneously ablating *ex vivo* tissue using an embedded $CO_2$ laser fiber and collecting the generated aerosol using silicone tubing for analysis by the REIMS system. The fiberbot is 1.65 mm in outer diameter and 10 cm in length.

**Movie S8.** Tele-micromanipulation of a selectively actuated fiberbot to reach four targets in sequence. The selectively actuated fiberbot is 2.00 mm in outer diameter and 1.2 m in total length. The selectively actuated section of the fiberbot is 12 cm in length.

**Movie S9.** Integration of the selectively actuated fiberbot with the DaVinci® robotic platform to simulate high precision tissue scanning along the contour of simulated tumor tissue.

**Movie S10.** *In vivo* demonstration of the laparoscopic robotic instrument in a porcine model. This movie demonstrates the fiberbot's simultaneous actuation and ablation (during the safe apnea phases) of spiral and circular paths in the liver, and a circular path in the cecum.
